# Subgenual Anterior Cingulate Cortex Functional Connectivity Abnormalities in Depression: Insights from Brain Imaging Big Data and Precision-Guided Personalized Intervention via Transcranial Magnetic Stimulation

**DOI:** 10.1101/2023.03.09.531726

**Authors:** Xiao Chen, Bin Lu, Yu-Wei Wang, Xue-Ying Li, Zi-Han Wang, Hui-Xian Li, Yi-Fan Liao, Daniel M. Blumberger, Francisco Xavier Castellanos, Eduardo A. Garza-Villarreal, Li-Ping Cao, Guan-Mao Chen, Jian-Shan Chen, Tao Chen, Tao-Lin Chen, Yan-Rong Chen, Yu-Qi Cheng, Zhao-Song Chu, Shi-Xian Cui, Xi-Long Cui, Zhao-Yu Deng, Qing-Lin Gao, Qi-Yong Gong, Wen-Bin Guo, Can-Can He, Zheng-Jia-Yi Hu, Qian Huang, Xin-Lei Ji, Feng-Nan Jia, Li Kuang, Bao-Juan Li, Feng Li, Tao Li, Xue Li, Tao Lian, Xiao-Yun Liu, Yan-Song Liu, Zhe-Ning Liu, Yi-Cheng Long, Jian-Ping Lu, Jiang Qiu, Xiao-Xiao Shan, Tian-Mei Si, Peng-Feng Sun, Chuan-Yue Wang, Han-Lin Wang, Xiang Wang, Ying Wang, Chen-Nan Wu, Xiao-Ping Wu, Xin-Ran Wu, Yan-Kun Wu, Chun-Ming Xie, Guang-Rong Xie, Peng Xie, Xiu-Feng Xu, Zhen-Peng Xue, Hong Yang, Jian Yang, Hua Yu, Yong-Qiang Yu, Min-Lan Yuan, Yong-Gui Yuan, Yu-Feng Zang, Ai-Xia Zhang, Ke-Rang Zhang, Wei Zhang, Zi-Jing Zhang, Jing-Ping Zhao, Jia-Jia Zhu, Xi-Nian Zuo, the DIRECT Consortium, Hua-Ning Wang, Chao-Gan Yan

## Abstract

**Background:** The subgenual anterior cingulate cortex (sgACC) plays a central role in the pathophysiology of major depressive disorder (MDD), and its functional interactive profile with the left dorsal lateral prefrontal cortex (DLPFC) is associated with transcranial magnetic stimulation (TMS) treatment outcomes. Nevertheless, previous research on sgACC functional connectivity (FC) in MDD has yielded inconsistent results, partly due to small sample sizes and limited statistical power. Furthermore, calculating sgACC-FC to target TMS individually is challenging.

**Methods:** Leveraging a large multi-site cross-sectional sample (1660 MDD patients vs. 1341 healthy controls) from Phase II of the Depression Imaging REsearch ConsorTium (DIRECT), we systematically delineated case-control difference maps of sgACC-FC. Then, we explored the potential impact of such group-level abnormality profiles on the TMS target localization and clinical efficacy. Next, we developed an MDD big data-guided individualized TMS targeting algorithm to integrate group-level statistical maps with individual-level brain activity to localize TMS targets individually.

**Results:** We found an enhanced sgACC-DLPFC FC in MDD patients compared to healthy controls (HC). Such group differences altered the position of the sgACC anti-correlation peak in the left DLPFC. In two independent clinical samples, we showed that the magnitude of TMS targets’ case-control differences in sgACC FC was related to clinical improvement. The MDD big data-guided individualized TMS targeting algorithm may generate individualized TMS targets that are clinically superior to group-level targets.

**Interpretation:** We reliably delineated MDD-related abnormalities of sgACC-FC profiles in a large, independently ascertained sample and demonstrated the potential impact of such case-control differences on FC-guided localization of TMS targets.

**Funding:** Ministry of Science and Technology of the People’s Republic of China, National Natural Science Foundation of China, and Chinese Academy of Sciences

## Introduction

Major depressive disorder (MDD) is a common and debilitating psychiatric disorder projected to be the most burdensome condition worldwide by 2030 ^1^. Despite extensive research, the pathophysiology of MDD remains elusive. Nevertheless, a key putative brain region or network hub appears to be the subgenual anterior cingulate cortex (sgACC), which shows reproducible metabolic hyperactivity ^2^, has been implicated in emotional responses, motivation, and rumination in MDD ^3^, and it has been shown to be an important target in deep brain stimulation and transcranial magnetic stimulation (TMS) ^4,5^. Repetitive TMS above 5 Hz on the left dorsolateral prefrontal cortex (DLPFC) indirectly stimulates the sgACC, and the closer to the sgACC target, the better the clinical outcome ^6^. Accordingly, identifying an optimized neuromodulation target in the left DLPFC based on sgACC-related functional connectivity (FC) is crucial for developing effective depression treatments ^7,8^. In light of inconsistent findings derived from studies with small sample sizes ^9–15^, we set out to establish a large sample to identify a reliable abnormal sgACC-DLPFC FC profile in MDD and further integrate this profile with individual brain activity to generate individualized neuromodulation targets for treating depression.

Numerous investigations have delved into FC abnormalities in MDD using resting-state functional magnetic resonance imaging (R-fMRI). Abnormal FCs between sgACC and amygdala, thalamus, temporal gyrus, lingual gyrus, cerebellum, DLPFC, and default mode network (DMN) regions such as medial and dorsal medial prefrontal cortex, precuneus, and parahippocampus have been reported ^9,10,13–20^. However, findings have been inconsistent, making integrating findings and generating precise profiles of sgACC-related FC abnormalities challenging. This deficiency in reproducibility could be partially due to small sample sizes, differences in preprocessing pipelines, and low statistical power of clinical imaging studies ^21,22^. To address the issue of limited sample size, we initiated the Depression Imaging REsearch ConsorTium (DIRECT) ^23^ and conducted an initial meta/mega-analysis (N_MDD_ = 1300), referred to as REST-meta-MDD ^24^. DIRECT Phase I shared ROI-level signals, thus enabling the investigation of multiple MDD-related abnormalities in network FC, FC topological and dynamic features, and functional lateralization ^24–31^. In DIRECT Phase II data reporting, we pooled an expanded MDD sample (N_MDD_ = 1660), which was preprocessed with a surface-based pipeline, DPABISurf ^32^. DIRECT Phase II shared voxel/vertex level BOLD time series, allowing more flexible and thorough investigations. Leveraging the most comprehensive MDD R-fMRI dataset to date encompassing depression patients and healthy controls, we can determine an aberrant sgACC-FC profile associated with MDD, characterized by superior reproducibility and low risk of false positives.

Maps of sgACC-related FC abnormalities are clinically useful for predicting repetitive TMS (rTMS) treatment outcomes in MDD patients ^33–35^. Specifically, the anti-correlation between sgACC and left DLPFC has been associated with clinical improvement from rTMS treatment ^36–40^. This has led to the intriguing notion that the FC between sgACC and left DLPFC could be leveraged to identify more precise rTMS targets and improve the efficacy of rTMS delivered to the left DLPFC ^7,41^. Researchers have identified a group-wise TMS target ^38^, which was the most anticorrelated DLPFC site to sgACC in the mean FC map from a large cohort of healthy adults. Subtle but significant case-control differences in resting-state FC profiles have been identified in a large sample of MDD patients ^24^. Thus, understanding the profiles of sgACC FC case-control differences and their impact on potential targets for rTMS applied to the left DLPFC could be a critical step toward developing optimized rTMS target site identification methods.

Individual human brains exhibit highly heterogeneous functional organization ^42^, with the DLPFC regions exhibiting the highest level of interindividual variation in cytoarchitecture, brain function, and network connectivity profiles ^8,43^. While several individualized FC-guided TMS target identification algorithms have been proposed ^36,41,44–48^, the target localizations of most existing TMS protocols have not been individualized. The major obstacles to identifying individualized TMS locations are the low signal-to-noise ratio in the sgACC area and the poor reproducibility of individual FC maps ^49,50^. The high reliability and statistical power of the DIRECT MDD cohort (N_MDD_ = 1660) allow the integration of group-level statistical maps and individual functional brain images to achieve precise and reliable TMS localization. Here, we propose an MDD big data-guided individualized TMS targeting algorithm based on dual regression (DR), which was initially developed for mapping group-level independent component analysis (ICA) results onto individual brains ^51^. During individualized target localization, the DR calculation is entirely confined to the DLPFC region, which has a high signal-to-noise ratio, avoiding noisy signals from the sgACC region. Thus, this approach enhances the efficacy and reliability of individualization approaches for identifying TMS targets ^52^.

In the present study, we leverage a large-scale multi-center sample (DIRECT Phase II, 1660 MDD patients and 1341 healthy controls (HCs)) to derive a reliable sgACC-related FC abnormality profile for MDD. Next, we showed that such case-control difference profiles may be related to the clinical efficiency of TMS and that the positions of the sgACC anti-correlation peaks might be different in the MDD patients as compared to the HCs. In light of this, we developed an MDD big data-guided individualized TMS targeting algorithm that may boost the clinical efficiency of TMS. We hypothesized that MDD patients would show a significantly abnormal sgACC-FC profile, especially in the left DLPFC. We also hypothesized that our newly developed DR-based approach would outperform traditional TMS group targets. To our knowledge, this is the first study to show the possible implications of the case-control abnormalities regarding the sgACC-FC profiles on the TMS target localization and to integrate large-scale group-level statistical maps with individual-level spontaneous brain activity to achieve individualized TMS targeting in MDD.

## Materials and methods

### Study sample

This study utilized four independent datasets. The first dataset (“DIRECT”) is a large-scale, multi-site consortium sharing standardized preprocessed R-fMRI time series. Building on the initial success of DIRECT Phase I (the REST-meta-MDD Project) ^24^, consortium members and international collaborators met on May 11^th^-12^th^, 2019, and agreed to launch DIRECT Phase II, which comprises 23 case-control designed datasets, including R-fMRI and T1 structural scans from 1660 MDD patients and 1341 HCs. Researchers from each site took a 2-day DPABISurf training course on September 14^th^-15^th^, 2019, to harmonize the organization and preprocessing of R-fMRI/T1 structural data. Demographic and clinical characteristics for each sample are presented in Figure 1 and Table 1. Site information, sample size, and previous publications based on the shared data are listed in Table S1. All participants were asked to self-report their sex (biological attribute) as part of the case report form (CRF). All participants in DIRECT Phase II were East Asian. Patients were diagnosed with MDD based on ICD 10 or DSM-IV. Healthy controls matched with MDD patients by age, sex ratio, and educational levels were recruited at each site. All participants provided written informed consent, and local institutional review boards approved each study from all included cohorts. The analysis plan of the current study has been reviewed and approved by the Institutional Review Board of the Institute of Psychology, Chinese Academy of Sciences (No. H21102). Data will be made available to the public as outlined in the Data Sharing Statement.

**Figure 1.**
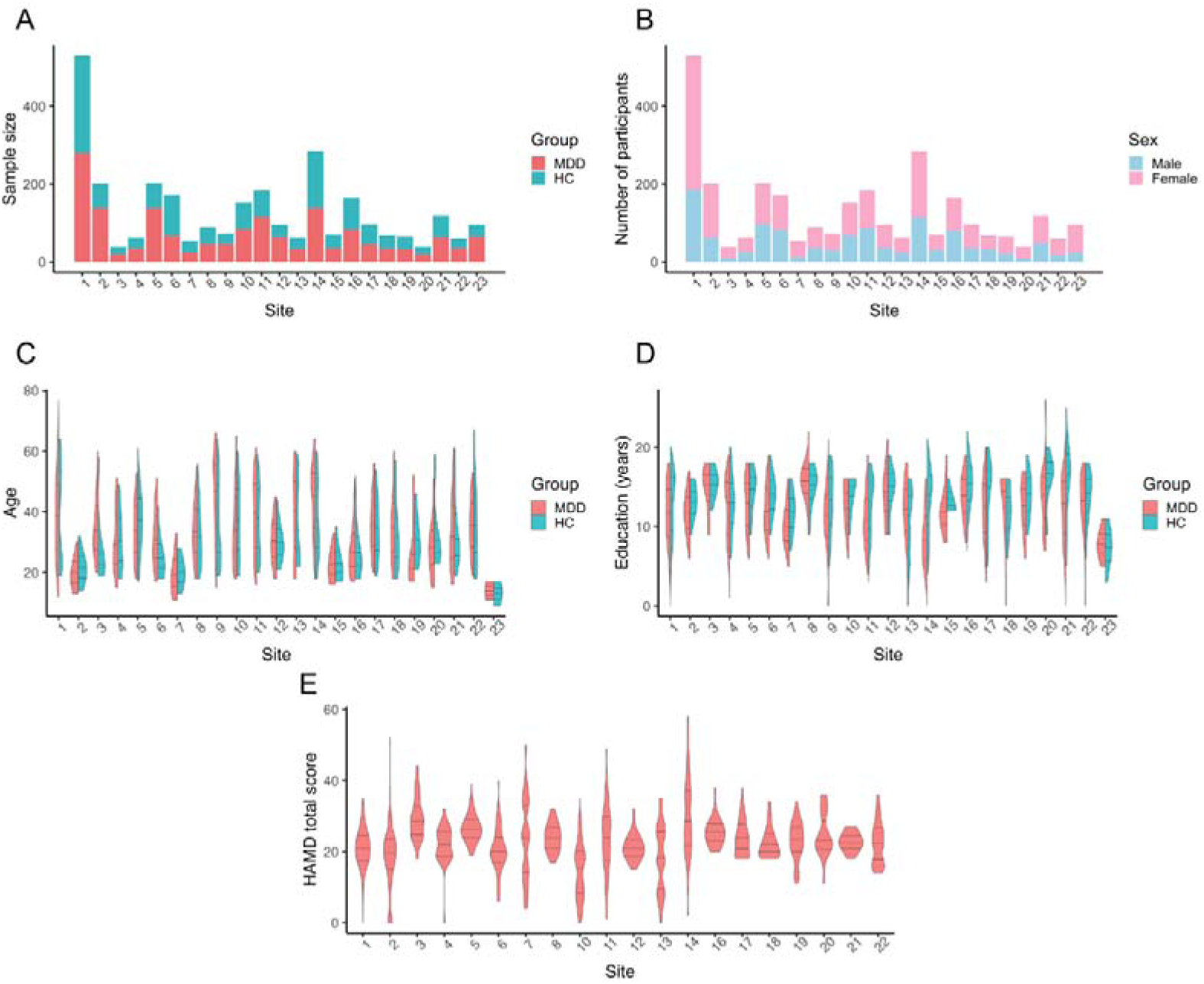
Sample characteristics of the DIRECT dataset. (A) Sample sizes of each site; (B) Number of male/female subjects irrespective of diagnosis; (C) Violin plots depicting the age distribution (in years). Solid black lines indicate the mean, 25^th^, and 75^th^ percentiles; (D) Violin plots show education distribution (in years). Solid black lines indicate the mean, 25^th^, and 75^th^ percentiles; (E) Violin plots depicting the distribution of scores of the Hamilton Depression Rating Scale (HAMD). Solid black lines indicate the mean, 25^th^, and 75^th^ percentiles.

**Table 1.**
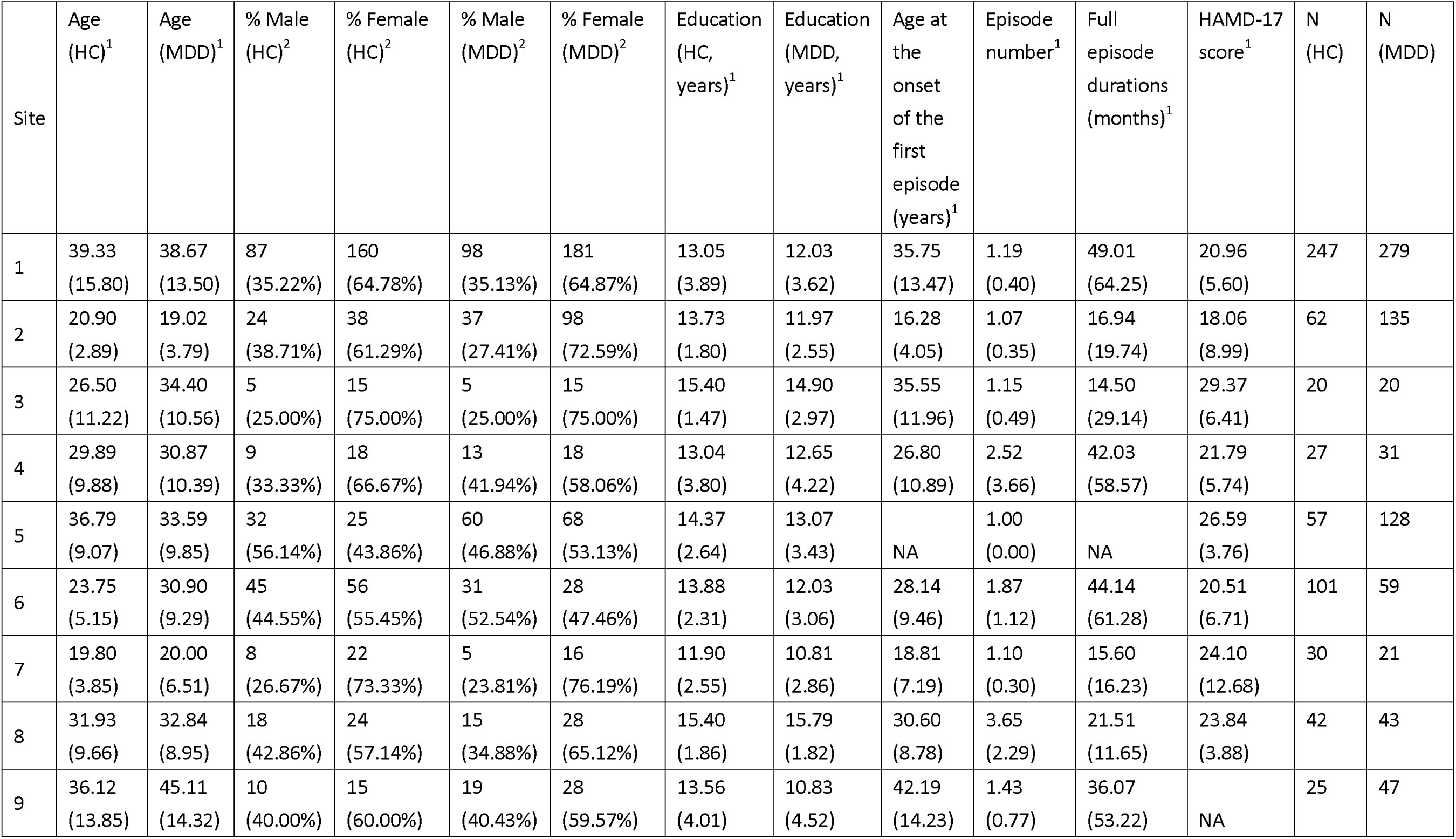
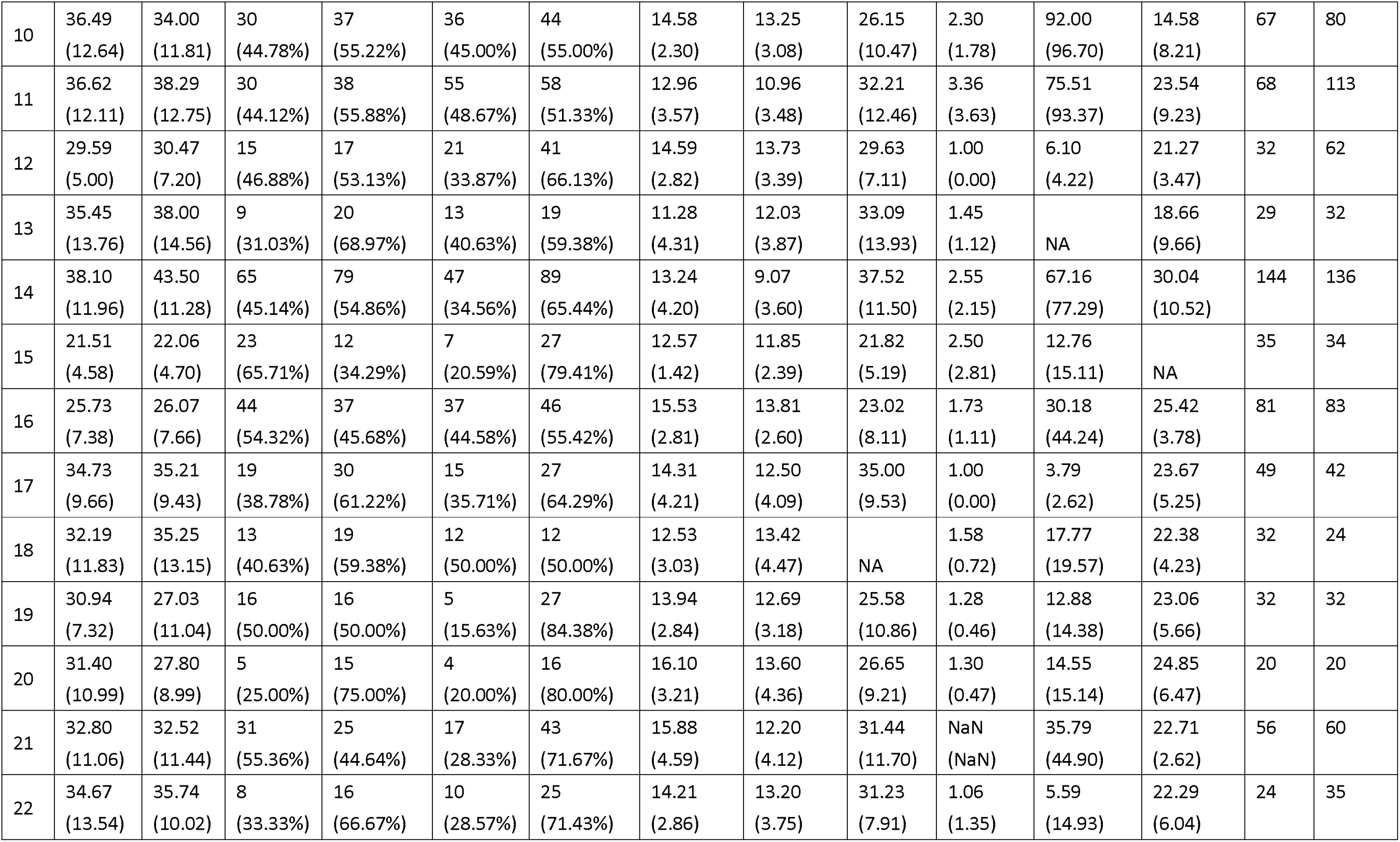
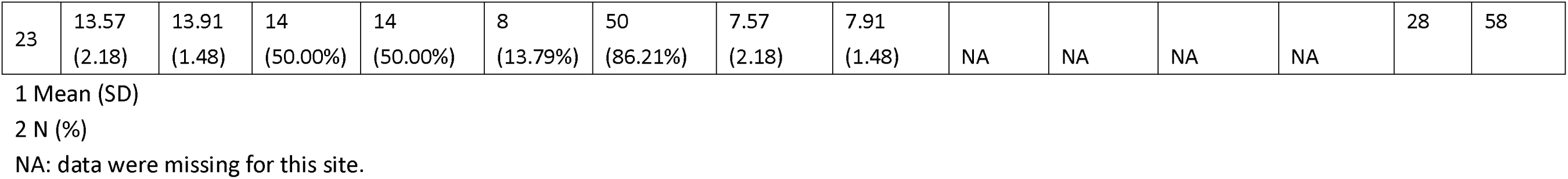
Demographic and clinical data for all samples included in the DIRECT II project.

The second dataset (“TRD-TMS”) comprises 25 medication-treatment-resistant MDD (TRD) patients who underwent 4 to 7 weeks of daily repetitive TMS applied over the left DLPFC. Patients’ TMS sites were recorded using their structural MRI images and a frameless neuronavigation system. Treatment response was assessed with the 24-item Hamilton Depression Rating Scale (HAMD). The targets for rTMS stimulation were determined using the 5.5-cm method. Only TMS outcomes and target coordinates were openly shared for this dataset. We obtained access to these data from the supplementary materials of Weigand et al. ^38^. For more details on this dataset, please refer to Weigand et al. ^38^.

The third dataset (“SID-TMS”) consists of 28 MDD patients with suicidal ideation who underwent 10 daily sessions of rTMS over the left DLPFC for 5 consecutive days. Clinical efficacy was evaluated using a 17-item HAMD. The TMS outcomes, neuroimaging data, participants’ demographic information, and target coordinates for this dataset were made available upon request, enabling evaluation of the performance of the MDD big data-guided individualized TMS targeting algorithm. The targets for individualized rTMS stimulation were determined by identifying the peak subunits in the DLPFC area with the most negative connections to the sgACC area in the original study. For more details on this dataset, see Li et al. ^53^.

The fourth dataset (“CUD-TMS”) comprises 27 cocaine use disorder (CUD) patients who underwent two daily sessions of rTMS treatment over the left DLPFC in an acute phase and two weekly sessions of rTMS treatment in a maintenance phase. The rTMS treatment was delivered at the left DLPFC using either the 5.5 cm anatomic criterion or the Beam F3 method. Depressive symptoms were a secondary treatment outcome in the original study. A subsample of 16 individuals, all with baseline HAMD scores above 7, was used for further calculation of individualized TMS targets. The TMS outcomes, neuroimaging data, participants’ demographic info, and target coordinates for this dataset were openly shared (https://openneuro.org/datasets/ds003037/versions/1.0.0). For more details on this dataset, see Garza-Villarreal et al. ^54^.

### Approach

The study’s first objective was to delineate case-control differences in the sgACC-FC profile and explore its implication in identifying FC-guided individualized TMS targets. Accordingly, we conducted a generalized linear model (GLM) to compare voxel-wise sgACC-FC maps of MDD patients and HCs in the DIRECT dataset. We then demonstrated the association between clinical improvement and group differences in TMS targeting sgACC-FCs by leveraging the TRD-TMS and SID-TMS datasets. Given that the peak sgACC anticorrelation of a normative connectome within the left DLPFC was usually selected as the FC-guided sgACC group target, we showed the impact of case-control differences on such group targets by separately identifying the peak sgACC anticorrelation in the mean sgACC-FC maps of the MDD group and HC group from the DIRECT dataset. Finally, we identified individualized optimal targets using the MDD big data-guided individualized TMS targeting algorithm guided by statistical maps (e.g., group difference map, mean sgACC-FC maps). We validated the clinical effectiveness of the individualized approach by computing the correlation between clinical outcomes and the distance between the actual TMS sites and the identified individualized targets in the SID-TMS and CUD-TMS datasets. All statistical tests conducted in the current study were two-sided.

### Power calculations for primary hypotheses

The primary outcome of the current study is the case-control differences regarding the sgACC-FC profiles. Estimates of the effect size (Cohen’s *d* = 0.186) of MDD patients’ abnormalities in FCs are drawn directly from our prior research based on the DIRECT Phase I dataset ^24^. Power calculation was performed using R version 4.3.1 ^55^ with pwr ^56^. A sample of 455 patients will achieve 80% power with a 5% Type I error rate.

### Image preprocessing

Acquisition parameters and scanners for all cohorts are provided in Table S2. All R-fMRI and structural MRI scans were preprocessed at each site using the same DPABISurf protocol, an R-fMRI data analysis toolbox evolved from DPABI/DPARSF ^32,57,58^ (For details, see SI). Given the controversy regarding global signal regression (GSR) and its essential role in identifying TMS targets ^41^, we performed preprocessing pipelines with and without GSR.

### FC maps of sgACC

Although recent studies have attempted to identify personalized TMS targets using surface-based algorithms ^44^, most previous studies have reported targets in volume-based MNI space with sgACC ROIs defined as a sphere in volume-based space ^36,59^. As a result, we used the volume-based preprocessed imaging data from DPABISurf to better compare our results with the existing literature.

We defined the sgACC as a 10 mm diameter sphere located on the average MNI coordinates based on prior studies showing reduced glucose metabolism or blood flow after receiving an antidepressant treatment (MNI coordinates: x = 6, y = 16, z = –10. For details, please refer to Fox et al., 2012 ^36^). The sgACC time series were determined for each individual by spatially averaging the preprocessed R-fMRI time series across all voxels in the abovementioned masks. We then calculated whole-brain FC maps in volume-based MNI space. FC was calculated using Pearson’s correlation and underwent Fisher’s r-to-z transformation. All FC maps were smoothed with a 6 mm full-width half maximum (FWHM) kernel size. We used ComBat ^60^ to control potential site and scanner biases (For details, see SI).

### Group difference maps of sgACC-FC profiles

We used a voxel-wise GLM to examine differences in the FC maps of sgACC between MDD patients and HCs in DIRECT Phase II. Cohen’s *f*^2^ was calculated to characterize the effect sizes of this group difference effect. The GLM model includes age, sex, education, and head motion as covariates:

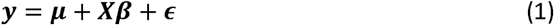

where y denotes the FC value of a given voxel from a given participant; **μ** stands for the constant term; X represents the design matrix for the covariates of interest (diagnosis, age, sex, education, and head motion); **β** is a vector of regression coefficients corresponding to X; and is a vector of residuals that follow *N*(0, **σ**^2^). Multiple comparison correction was conducted using false discovery rate (FDR) correction at *q* < 0.05.

To further interpret group difference maps, we extracted the mean FC values of seven networks using Schaefer’s 400 parcellation atlas ^61^. A GLM model identical to model (1) was constructed to characterize case-control differences for each network. Bonferroni multiple comparison correction was conducted (*p* < 0.05/7). We further explored the effect of the identified DLPFC clusters in several subgroups. Specifically, patients who were in their first episode and had never received any antidepressant medication treatment (first episode drug naïve, FEDN, N = 484) and patients who had undergone more than one episode (recurrent, N = 439) were selected and compared. Three contrasts, FEDN vs. HC, recurrent vs. HC, and FEDN vs. recurrent, were analyzed.

### Relationship between group differences in TMS targets’ sgACC FCs and clinical outcomes

To explore the relationship between group differences in TMS targets’ sgACC FCs and clinical outcomes, we first extracted the mean *t*-values from the group difference map of 8 mm radius spheres centered at each targeting coordinate in the TRD-TMS dataset, then examined the Pearson correlations between these *t*-values and HAMD score reductions. We anticipated that greater group differences in sgACC-FC at the target location (i.e., higher *t* values) would be related to better TMS therapeutic effects (higher HAMD reductions). To test the robustness of our findings, we also used spheres with 2 mm, 4 mm, and 10 mm radiuses to extract the *t*-values of group differences.

### Group targets based on mean sgACC-FC maps

The prior group-level DLPFC TMS target had been derived from a cohort of healthy young adults ^38^; here, we separately averaged whole-brain sgACC-FC maps across all the DIRECT participants in the MDD and HC groups. We then searched for the peak sgACC anticorrelated voxel within the DLPFC area (i.e., Brodmann area (BA) 46) as the mean sgACC-FC guided TMS targets for the MDD and HC groups.

### Identification of individualized TMS targets

The reliable statistical maps from the DIRECT big sample best reflect the probability of MDD-related abnormalities in sgACC-FC. Therefore, we can use these maps to guide the identification of individualized abnormalities by combining this big-data-based abnormality information with the individualized R-fMRI data from a given patient, obtaining reliable statistical maps from the DIRECT sample. We used the dual regression approach to identify individualized TMS targets guided by group-level statistical maps in the SID-TMS and CUD-TMS datasets. Dual regression is a common method in independent component analysis (ICA) for projecting group-level independent components (e.g., functional networks) onto the individual subject level (see Figure S4 for details). In the first step of the MDD big data-guided individualized TMS targeting algorithm, a group-level statistical spatial map (e.g., the sgACC-FC group difference map reflecting the probability of MDD-related abnormalities in sgACC-FC) was used as a spatial regressor in the GLM to identify the temporal dynamic of the group-level map (similar to spatial correlation with the abnormality spatial map). A time series associated with the spatial map of MDD-related FC abnormalities was generated. In the second step, the derived time series was used as a temporal regressor in the GLM to identify an individual-level spatial map (similar to the temporal correlation with the previous time series). This spatial map can be considered the best-individualized abnormality guided by big-data-based abnormality. Given our prior knowledge of DLPFC TMS treatment in MDD, we confined the big-data-based abnormality dual regression to the DLPFC area. That is, we use the group DLPFC abnormality probability map to find the individualized DLPFC target in a given MDD patient. The final coordinates for the individualized TMS targets are defined as the centroids of the largest clusters within this DLPFC region on the individual-level spatial maps. Additionally, we calculated the individualized target coordinates using the seed map approach, following the methods described by Fox et al. ^41^ and Cash et al. ^44^. In the seed-map approach, a seed time series is extracted by computing a weighted average time series of all voxels within the seed map (e.g., a group average map of sgACC-FC, but excluding the DLPFC area). Subsequently, Pearson’s correlation coefficients are computed between this extracted time series and all other DLPFC voxels. The final TMS target is the most negatively functionally connected cluster in the DLPFC area. Of note, in the seed map approach, the goal of the first step is to find the most sgACC-like time series, which is not confined to the noisy sgACC area. Following this rationale, the DLFPC time series should not be included to avoid biasing the estimation of the sgACC-like time series. Thus, the DLPFC area was excluded. Thus, the exclusion and inclusion of DLPFC differs between the seed map approach and the DR approach due to the different underlying rationales. Details of the individualized TMS target localization algorithms are provided in the supplementary materials.

### Clinical efficacy of the group-level and individualized TMS targets

We leveraged the SID-TMS and CUD-TMS datasets to evaluate the clinical significance of individualized TMS targets. We identified the proposed individualized TMS targets from the MDD big data-guided individualized TMS targeting algorithm and calculated the targeting offset (i.e., Euclidean distance between the individualized optimal TMS targets and the actual stimulation coordinates) for each patient. Subsequently, we calculated the Pearson correlations between clinical improvement (i.e., HAMD reductions) and targeting offset. We anticipated a negative correlation between clinical outcomes and target offset (i.e., the closer the actual stimulation target was to the individualized target from the MDD big data-guided individualized TMS targeting algorithm, the higher the clinical improvement). Age, sex, and head motion were included as covariates in the regression models when calculating correlations between targeting offsets and clinical improvement.

## Results

### Group difference maps of sgACC-FC

In the large-scale DIRECT Phase II dataset, we found significant MDD-related hyperconnectivity with the sgACC in bilateral DLPFC, temporal parietal junction, and occipital lobe, as well as hypoconnectivity in the bilateral temporal lobe, left inferior frontal gyrus, and left postcentral gyrus when preprocessing included GSR (Figure 2A). When GSR was not included in preprocessing, MDD-related sgACC FC alterations showed predominantly hypoconnectivity. Such abnormally decreased FCs were found across the central gyrus, occipital lobe, insular cortex, temporal lobe, and a small portion of the frontal lobe. Without GSR, MDD-related hyperconnectivity was limited to subcortical regions (Figure S1A). Given that significant case-control differences in the DLPFC area were revealed only when GSR was implemented, subsequent analyses were based on results with GSR. The uncorrected group difference maps calculated in the volume space showed remarkable similarity with those in the surface space (Figure S2). Network-wise FC analyses showed that MDD patients’ FC between sgACC and the limbic network (LN) was significantly reduced compared to HC (*t*(2880) = –4.122, *p_corrected_* < 0.001, Cohen’s *d* = 0.171). The FC between sgACC and the frontoparietal network (FPN) was enhanced and approached significance (*t*(2880) = 2.419, *p_corrected_* = 0.055, Cohen’s *d* = 0.090, Figure 2B). Without GSR, MDD patients showed decreased FC between sgACC and all brain networks (all *p_corrected_* < 0.05) except for the FPN (Figure S1B). We identified two contiguous clusters of voxels that showed significant group differences in the left DLPFC. Group difference cluster 1 (MNI coordinates: x = –44, y = 38, z = 32; *t*(2880) = 3.277, *p* < 0.001, Cohen’s *d* = 0.141) was ventral to group difference cluster 2 (MNI coordinates: x = –34, y = 36, z = 40; *t*(2880) = 3.670, *p* < 0.001, Cohen’s *d* = 0.126) (Figure 2C). In subgroup analyses, when GSR was performed, FEDN patients showed enhanced FCs in both clusters (cluster 1: *t*(1790) = 2.282, *p* = 0.023, Cohen’s *d* = 0.124; cluster 2: *t*(1790) = 2.273, *p* = 0.023, Cohen’s *d* = 0.123). There was a significant enhancement in the DLPFC cluster 2 between the recurrent MDD patients and HCs (*t*(1745) = 2.765, *p* = 0.006, Cohen’s *d* = 0.159) while cluster 1 approached significance (*t*(1745) = 1.864, *p* = 0.063, Cohen’s *d* = 0.107). No significant difference was revealed between the FEDN and recurrent patients (see Figure 3).

**Figure 2.**
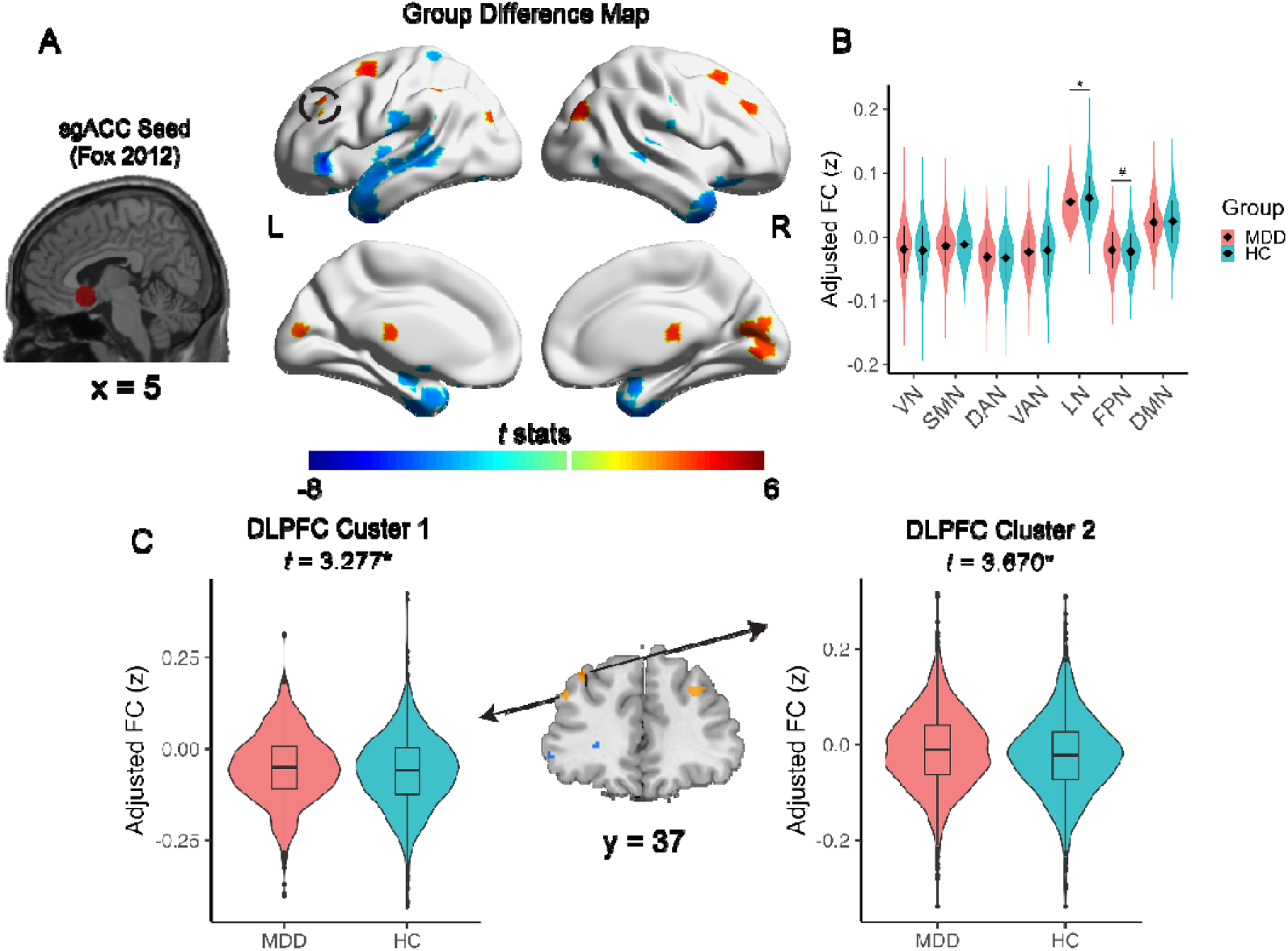
Group differences of subgenual anterior cingulate cortex (sgACC) functional connectivity (FC) profiles are related to TMS treatment efficacy, demonstrating clinical significance. (A) Two-sample t-test maps of MDD-related sgACC FC abnormalities with global signal regression (GSR) implemented. (B) Group differences of FCs between sgACC and visual network (VN), somatomotor network (SMN), dorsal attention network (DAN), ventral attention network (VAN), limbic network (LN), frontoparietal network (FPN), and default mode network (DMN). (C) Two clusters showed significant case-control differences in sgACC-FC. Abbreviations: DLPFC, dorsal lateral prefrontal cortex; L, left hemisphere; R, right hemisphere. *: significant after Bonferroni correction; #: approaching significance.

**Figure 3.**
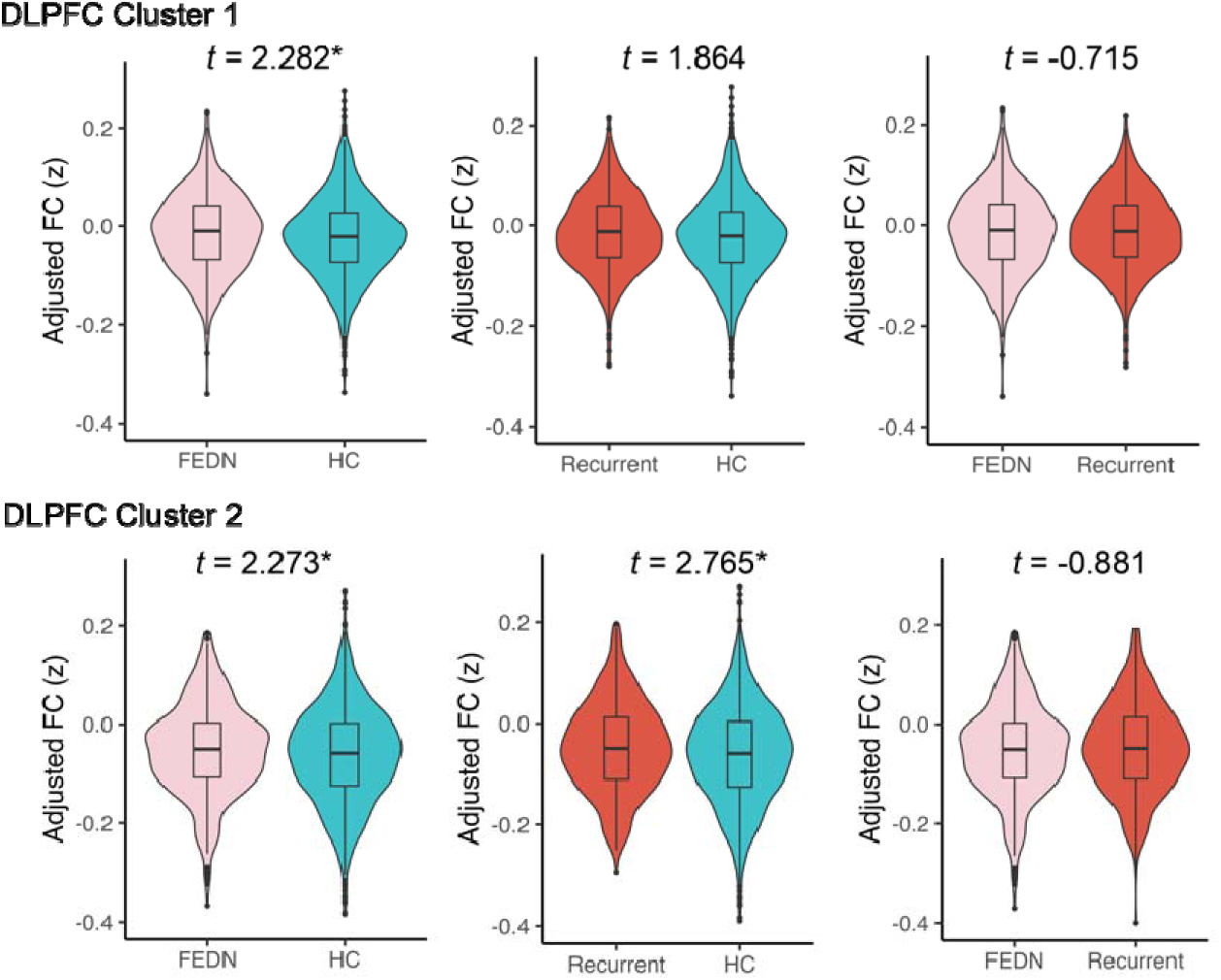
Subgroup differences regarding two clusters found in DLPFC. Abbreviations: FEDN, first episode drug naïve; HC, healthy control.

### sgACC anticorrelation peaks in MDD and HCs

Our results highlighted the case-control differences in sgACC-FC profiles. Since the prior sgACC group target (MNI coordinates: x = −42, y = 44, z = 30) had been based on a cohort of young, healthy adults^38^, we sought to examine potential differences in anticorrelation peaks extracted from the mean sgACC FC maps of MDD and HC groups (Figure 4B-C) in the DIRECT dataset. We found that the anticorrelation peak of MDD patients (MNI coordinates: x = –40, y = 50, z = 22) differed from that of HCs (MNI coordinates: x = –42, y = 38, z = 32), probably due to abnormal FCs within the left DLPFC in MDD patients. The anticorrelation peak extracted from the DIRECT HCs was closer to the previously reported locus^38^ (Figure 4).

**Figure 4.**
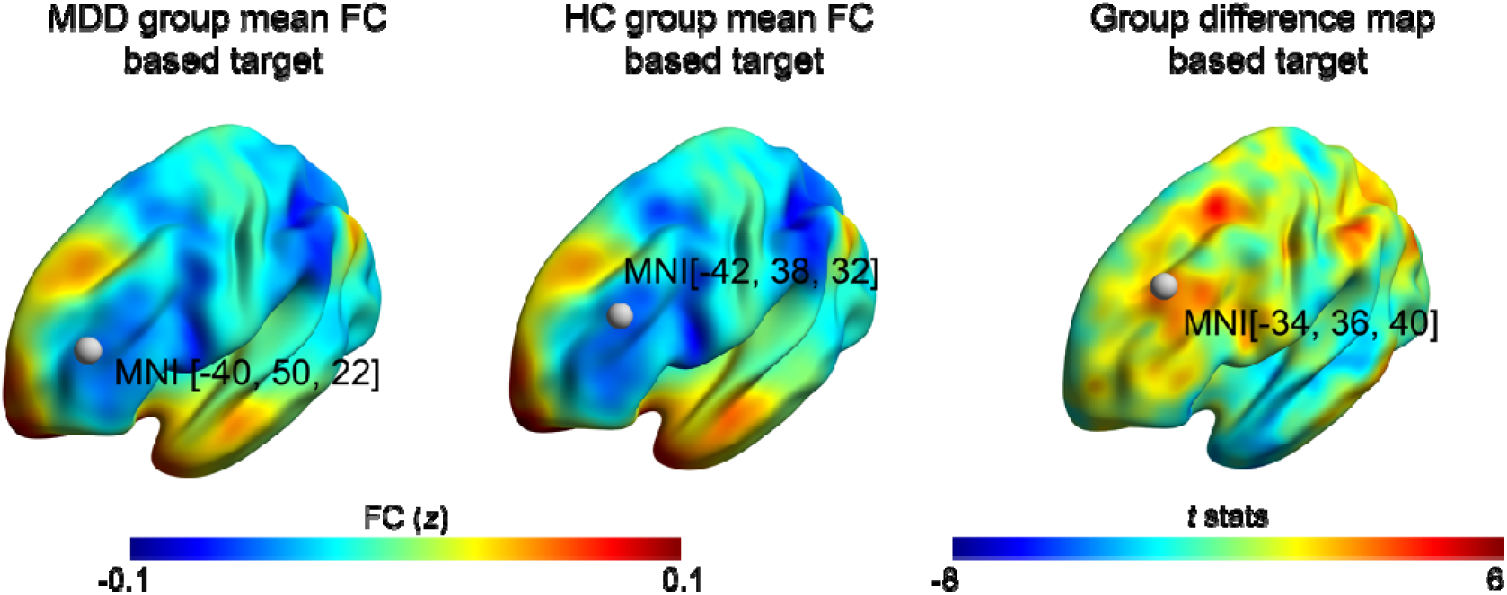
The peak of the sgACC anticorrelation in MDD patients differed from that in HCs.

### sgACC-FC group differences correlate with TMS treatment outcomes

Using the TRD-TMS dataset, we examined the relationship between group differences (t-values) and clinical outcomes (HAMD reductions) to test the clinical relevance of the group difference maps. Group differences were positively correlated with HAMD score reductions (r(23) = 0.448, p = 0.025, Figure 5A), suggesting that group-level difference maps may be useful for enhancing the outcomes of TMS for treating MDD by improving target localization. Significant associations were also observed using different radius settings (Figure S11). The same trend was observed in the SID-TMS dataset, albeit it failed to achieve significance (r(26) = 0.304, p = 0.108, Figure 5).

**Figure 5.**
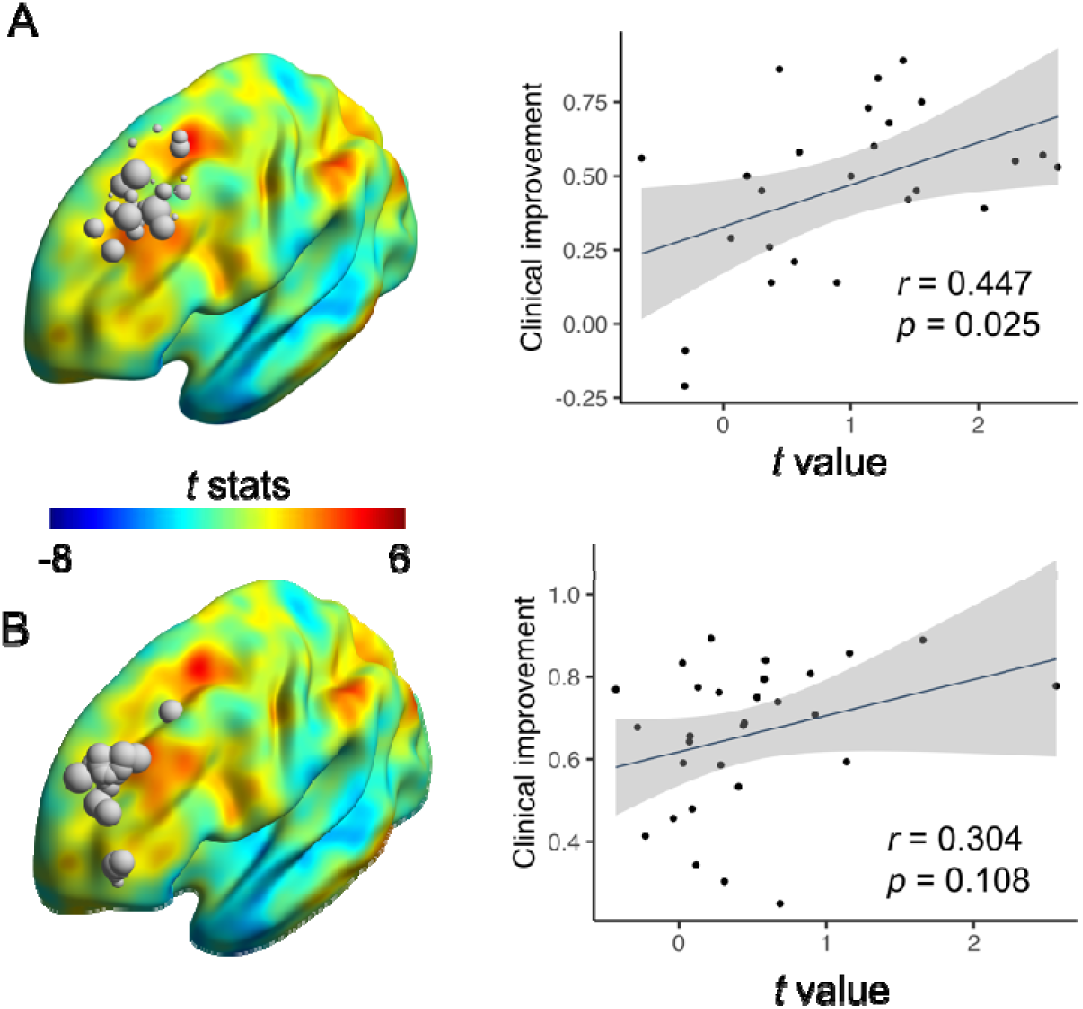
The group difference regarding sgACC-FC to the TMS targets was correlated with clinical efficacy. (A) The TMS targets were extracted from the TRD-TMS dataset. The sizes of the spheres indicate the magnitudes in Hamilton Depression Rating Scale (HAMD) reductions, with the group difference map rendered on the surface. The scatter plot depicted that the magnitudes of the group differences in the FC between TMS targets and sgACC were positively related to clinical improvements in the TRD dataset. (B) Findings were replicated in the SID-TMS dataset^53^. Abbreviations: FC, functional connectivity; TMS, transcranial magnetic stimulation.

### Individualized targets suggest higher clinical efficacy than group targets

We utilized the MDD big data-guided individualized TMS targeting algorithm to calculate individualized optimal targets for each SID-TMS and CUD-TMS dataset participant. Clinical relevance of the targets was determined by the correlation between target offset and clinical improvement. The individualized target locations derived from the sgACC-FC group difference map are illustrated in Figure 6. The individualized targets (SID-TMS: r(26) = –0.562, p = 0.002; CUD-TMS: r(14) = –0.511, p = 0.037) outperformed their corresponding group-level targets (SID-TMS: r(26) = –0.349, p = 0.044; CUD-TMS: r(14) = –0.167, p = 0.293) in both datasets (Figure 6). The DR-based individualized targets derived from the MDD or HC group-average sgACC-FC maps also outperformed their corresponding group-level targets and seed map-based targets (Figure S6-8). Among all the individualized targets, the DR-based targets guided by the group difference map achieved the highest clinical efficacy.

**Figure 6.**
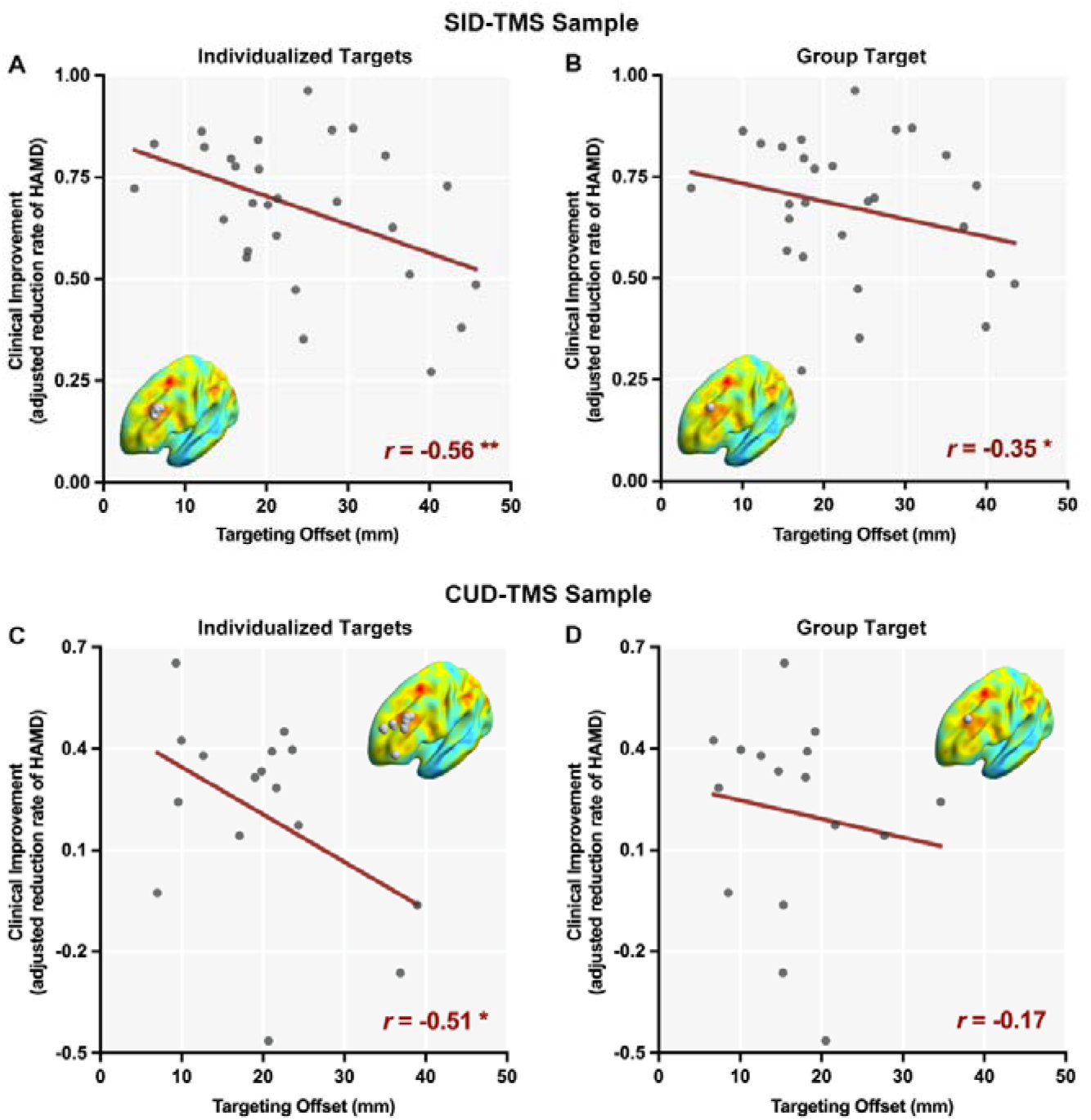
Individualized targets derived from the group difference map exhibited greater clinical efficacy than the corresponding group targets. Clinical efficacy was characterized by computing the correlations between the offset distances of TMS targets and the clinical improvements observed in the CUD-TMS dataset. The TMS targeting offset distance was defined as the Euclidean distance between the actual rTMS stimulation coordinates and the individualized or group targets. Clinical improvement was defined as the HAMD reduction during rTMS treatment, adjusted for age, sex, and head motion. The locations of the targets are displayed on the cortex. The sizes of the spheres indicate the magnitudes of Hamilton Depression Rating Scale (HAMD) reductions. (A-B) Clinical efficacy of the individualized and group targets in the SID-TMS sample. (C-D) Clinical efficacy of the individualized and group targets in the CUD-TMS sample. *p < 0.05. ** p < 0.01.

## Discussion

In the present study, we leveraged a large multi-site fMRI sample (1660 MDD patients and 1341 HCs) and three independent TMS datasets to delineate abnormalities in sgACC FC in MDD and explore their potential impact on the localization of TMS targets. Specifically, with GSR implemented, we found enhanced FCs between sgACC and left DLPFC, bilateral supplementary motor areas and inferior parietal lobes, thalamus, and visual areas, and decreased FCs between sgACC and left anterior insula, left superior temporal lobe, and bilateral temporal poles in MDD patients. Patients with MDD exhibited significantly reduced FC between the sgACC and the DMN, while FC between the sgACC and the FPN was only marginally increased. Crucially, we showed that the clinical outcomes of TMS treatments were related to the magnitude of the case-control differences in the FCs between sgACC and TMS targets. Furthermore, such group difference profiles altered the position of the sgACC anti-correlation peak in the left DLPFC. Additionally, the MDD big data-guided individualized TMS targeting algorithm to identify individualized TMS targets showed better clinical efficacy than TMS targets based on group sgACC-FC profiles.

### MDD-related FC abnormalities of sgACC

Our results add to a growing literature documenting functional network abnormalities involving the sgACC in MDD ^9,11–13,17,19,62–64^. Nevertheless, notable discrepancies in the type of abnormality (enhanced/reduced) and specific brain regions showing altered sgACC FCs have been reported. Considering the small effect sizes (Cohen’s *f*^2^ < 0.01) of MDD-related sgACC-FC abnormalities, the limited sample sizes in previous studies entail a high risk of false positive findings ^21,65,66^. Another potential source of heterogeneity in previous findings may be whether or not they applied GSR (See supplementary materials for detailed discussion).

With an unprecedented sample size, our results provide among the most robust evidence to date. Specifically, MDD patients showed enhanced sgACC-thalamus FC and decreased sgACC-limbic network FC regardless of whether GSR was implemented. These results are consistent with previous studies in adolescents ^19^ and adults ^59^ with MDD. Previous studies have reported abnormal FCs between sgACC and limbic areas and some subcortical regions, such as the amygdala ^9,20^ and parahippocampus regions ^13^. Together, the present results are consistent with a model highlighting sgACC as a critical hub in an “extended medial network,” which also encompasses limbic, thalamic, and striatal regions and plays a key role in the pathophysiology of MDD ^67,68^. This “extended medial network” overlaps substantially with the DMN. Indeed, decreased FCs between sgACC and DMN regions, such as the medial prefrontal cortex, precuneus, temporal gyrus, and parahippocampus regions, were revealed in MDD relative to HCs when GSR was not implemented. Such abnormalities have been previously reported ^10,12,15,34^, which led to the hypothesis that abnormally enhanced FC between sgACC and DMN are the network underpinnings of rumination ^69^. However, contrary to the aforementioned hypothesis, we found reduced, instead of enhanced FC between sgACC and DMN. Similarly, in the first phase of DIRECT, we demonstrated that MDD was characterized by reduced FC within DMN ^23,24^. We note that the first and second phases of the DIRECT data are solely comprised of Chinese samples, while most studies that have reported enhanced sgACC-DMN FCs have been in Caucasian samples. Different prevalence rates ^70,71^, heterogeneous symptoms ^72^, and different risk alleles ^73^ have been reported in Caucasian and Eastern Asian groups. Accordingly, we cannot exclude racial differences contributing to this discrepancy. It is worth noting that we found enhanced FCs between the visual region and sgACC in MDD patients relative to HCs when GSR was implemented, while significantly reduced sgACC-visual region FCs were revealed when GSR was not performed. Most of DIRECT II sites’ R-fMRI data were collected with participants’ eyes closed (22 out of 23 sites). Prior research had shown that participants are more likely to fall asleep when their eyes are closed during data acquisition, and drowsiness may alter FC patterns in visual regions ^74^. Thus, it is possible that abnormalities in visual region FCs may be due to MDD patients’ lower levels of wakefulness.

### Clinical relevance of abnormal sgACC-DLPFC anticorrelation in MDD

Once the group difference maps of sgACC FC profiles were delineated, we further explored the impact of such an abnormality, especially in DLPFC, on identifying TMS targets. Most clinical trials have focused on applying TMS to the left DLPFC based on the hypothesis that high-frequency rTMS will enhance hypoactivity during depressive episodes ^8,75^. The DLPFC is anatomically extensive ^76^. However, which DLPFC sub-field is the best target for TMS remains unclear. The current FDA-approved protocol (i.e., the “5 cm” method) leads to large interindividual variation in stimulation sites, which may contribute to the heterogeneity in the effect sizes of antidepressant responses in prior trials ^77,78^.

Previous targeting approaches leveraging anatomical landmarks has not consistently outperformed the “5 cm” method or the F3 Beam method ^79,80^. Considering the unsatisfactory effect of TMS target localization based on brain anatomical parcellation, group-level normative sgACC anticorrelation peaks based on healthy population datasets have been frequently used as TMS targets in recent years ^36,38,41^. However, in the present study, we found that the locations of such anticorrelated peaks differ substantially between MDD and HC samples when measured in a large clinical cohort. Therefore, it might be problematic to identify TMS targets based solely on sgACC-FC profiles in healthy or depressed individuals. Intriguingly, we found that the magnitudes of case-control differences in TMS targets’ FCs to sgACC were positively related to the clinical improvements after receiving rTMS to the left DLPFC. Such correlation implies that the case-control differences in the FC between sgACC and the left DLPFC might bear important information that could be leveraged to guide the identification of reliable, individualized TMS targets.

### The MDD big data-guided individualized TMS targeting algorithm may improve the clinical efficacy of TMS targets

In the current study, we developed an MDD big data-guided individualized TMS targeting algorithm to individualize the TMS targets derived from group-level statistical maps. The proposed approach takes advantage of the high signal-to-noise ratio and reliability of large sample statistical maps while integrating individual spontaneous brain activity of individuals with MDD. Most existing individualized TMS target localization algorithms are based on calculating sgACC FCs using densely sampled MRI images from single subjects ^81^. However, individual MRI images tend to be noisy and unreliable ^82,83^, especially in the sgACC region. Air in the sinuses often introduces susceptibility artifacts due to the different magnetic properties of air and brain tissue. Signal loss and geometric distortion are common in areas close to air-filled sinuses, such as the inferior frontal cortex, including the sgACC ^84^. The seed map approach has been proposed to alleviate such difficulties due to the subpar image quality of the sgACC region ^41,44^. In the seed map approach, all voxels within the seed map (except for the DLPFC region) were used to extract the seed time series to improve its signal-to-noise ratio. However, considering that the weight (e.g., FC value) of the sgACC area is usually extremely high in the seed map, the derived seed time series remains somewhat similar to the noisy sgACC time series and doesn’t achieve the best TMS localization. A cutting-edge MDD TMS therapy combining FC-guided target localization, high dose, and intermittent theta-burst stimulation (iTBS) was reported to be highly effective in a randomized, double-blinded, sham-controlled clinical trial ^45,46^. However, the targeting algorithm relied on hierarchical agglomerative clustering in the sgACC area which has a low signal-to-noise ratio. Nevertheless, the final target was still determined according to individual-level sgACC-DLPFC FCs. For the proposed MDD big data-guided individualized TMS targeting algorithm, we view the reliable statistical maps from the DIRECT big sample as the best reflection of the probability of MDD-related abnormalities in sgACC-FC. Therefore, we used these maps to guide the identification of individualized abnormalities by combining this big-data-based abnormality information with the individualized R-fMRI data from a given patient. Given the a priori knowledge of DLPFC TMS treatment in MDD, we confined the big-data-based abnormality dual regression only within the DLPFC area, which is less affected by susceptibility artifacts. In this way, the superior signal quality of DLPFC and the effective and reliable properties of the dual regression algorithm enhance the accuracy of target localization.

Encouragingly, the DR-based individualization targets enhanced the clinical significance of corresponding group-level targets, regardless of the template used. This result supports the generalizability and extensibility of the algorithm, offering the potential for TMS targeting based on other circuits and biomarkers. Considering most existing TMS research still relies on the traditional 5 cm or Beam F3 methods for targeting, one approach based on our data to improve targeting would be simply shifting the target to a more anterior and lateral position. However, such a simple shift was not supported by the present study. Instead, 25% of targets individualized using the group difference map were more medial than the original targets, and 50% of targets individualized by the group difference map were more posterior than the original targets. Therefore, the MDD big data-guided individualized TMS targeting algorithm does not simply set a more anterior and lateral coordinate in the BA46 area as the individualized optimization target for subjects. Among the group-level templates used in the MDD big data-guided individualized TMS targeting algorithm, the sgACC-FC group difference map performed the best, instead of the commonly used average sgACC based on healthy individuals. This may reflect the abnormal posterior shift of the MDD sgACC anticorrelation peak we found, and it emphasizes the immense clinical value of examining spontaneous brain activity differences between MDD and HC in a large sample. Previous studies either included only a small quantity of MDD functional MRI data ^46,47,85^ or developed targeting algorithms based on large-scale HC samples ^49^. In this study, we utilized an unprecedented amount of functional MRI data from MDD and HC, obtained a reliable group-level difference map of sgACC-FC, and successfully validated its potential in treating MDD with TMS.

### Limitations

Several limitations need to be considered. First, we noted that the inconsistency between our results and the broader literature could partly be due to racial differences of the samples (i.e., Eastern Asian vs. Caucasian). Efforts that intend to pool existing neuroimaging data worldwide, such as the Enhancing NeuroImaging Genetics through Meta-Analysis (ENIGMA) Major Depressive Disorder (MDD) consortium ^25^, have accumulated large-scale data primarily from Western countries and published several high-impact studies delineating MDD-related anatomical abnormalities ^86,87^. Planned collaborations between the DIRECT and the ENIGMA-MDD consortiums are ongoing to help address potential cultural, genetic, and environmental mechanisms in more diverse groups of MDD patients. Second, we performed volume-based preprocessing to facilitate comparison with the previous literature.

Surface-based preprocessing strategies have provided more accurate and detailed representations of cortical and subcortical structures ^88^. Recent research has begun to explore surface-based rTMS target identification algorithms and has shown promising clinical relevance ^39,40,89^. Future research should consider using surface-based case-control difference maps to further refine ways to identify TMS targets. Given the established involvement of the left DLPFC in existing TMS treatment protocols, the present study restricted investigation to within this area.

We noted that other brain regions (e.g., angular gyrus and supplementary motor area) showed significant case-control differences worth further research and may serve as potential targets for neuromodulation ^90,91^ ^92^. The MDD big data-guided individualized TMS targeting algorithm can be readily transferred to other neural circuits or other brain imaging-derived feature maps (e.g., ICA, functional gradient, normative modeling). The clinical efficacy of these alternative targets is worthy of future investigations. Identifying a reliable personalized TMS target solely based on an individual’s R-fMRI data (around 8 mins of fMRI scan in clinical practice) is challenging. Due to the poor replicability of FC ^93^, existing individual-level network parcellation algorithms need a large quantity of fMRI images (usually more than one hour of scanning time) ^94^. The present study utilized three independent TMS samples to validate the efficacy of the individualized algorithm. Nevertheless, the two TMS datasets used to validate the individualized TMS targets are limited in sample size ^37,95^. Publicly available TMS brain imaging datasets could be used for independent validation of target localization algorithms to reduce the false positive rate; however, access to such datasets remains challenging. In addition, prospective, double-blind clinical trials are warranted to compare the treatment outcome across different rTMS targeting algorithms (e.g., traditional anatomical landmark-based targeting, group-level sgACC-FC targeting, and MDD big data-guided individualized TMS targeting algorithm). Therefore, we call upon researchers involved in this field to publicly share data on TMS targets, clinical efficacy, and brain imaging, and we will also openly share data from our related prospective studies ^96^. The large sample size of the DIRECT consortium aggregated dataset allows for intriguing analyses, such as bio-subgroups of MDD patients. Indeed, some previous DIRECT studies have shown that MDD patients can be subgrouped ^30,97^. Future studies may further determine whether bio-types could be achieved using sgACC ^91^. Most of the DIRECT II R-fMRI data were acquired when participants were instructed to close their eyes, which has been shown to be associated with an increased likelihood of sleep during scanning ^74^. Instructing participants to keep their eyes open and look at fixation can help prevent participants from falling asleep and is easy to apply. Since large multi-site R-fMRI data aggregation endeavors such as DIRECT are prone to be biased by non-neurophysical factors such as head motion, sleepiness, etc. ^98^, it is important to prospectively apply well-designed standard operation procedures in future large-scale multi-site scientific projects.

## Conclusion

In summary, we leveraged a large sample of MDD patients to fully delineate group differences in sgACC-FC maps between MDD patients and HCs. We next demonstrated the impact of such case-control differences on group TMS targets based on sgACC-FC profiles by showing that the magnitudes of case-control differences in the FC between sgACC and TMS targets were positively associated with clinical outcomes and the peak sgACC anticorrelation locations were different in MDD patients as compared to HCs. Moreover, we developed an MDD big data-guided individualized TMS targeting algorithm to identify individualized TMS targets and demonstrated that this approach may improve clinical efficacy compared to group targets based on sgACC-FC profiles.

### Contributors

C-GY, Y-FZ, X-NZ, XC, and BL conceived and designed the study. C-GY, Y-FZ, J-PZ, and X-NZ coordinated the collaboration. JQ, LK, T-MS, TL, K-RZ, Z-NL, L-PC, JY, X-PW, Y-GY, C-YW, C-MX, G-RX, Y-SL, Y-QY, XW, YW, X-FX, Y-QC, Q-YG, W-BG, J-PZ, YH, H-NW, B-JL, WZ, and J-PL supervised data collection. X-RW, QH, Y-KW, HY, A-XZ, Y-CL, J-SC, P-FS, X-YL, FL, C-CH, X-LC, F-NJ, J-JZ, X-LJ, G-MC, Z-SC, T-LC, X-XS, TC, B-JL, M-LY, Z-PX and BL organized the data. XC and C-GY analyzed and interpreted the data. BL and C-GY designed the dual regression-based TMS targeting algorithm. H-NW and B-JL collected the SID-TMS dataset. XC, BL, and C-GY drafted the manuscript. All authors revised the manuscript for important intellectual content and approved the final submitted version. XC, BL, and C-GY accessed and verified the data. XC, BL, and C-GY had full access to all the data in the study. C-GY had final responsibility for the decision to submit for publication. All authors were responsible for the final decision to submit for publication and have seen and approved the manuscript.

### Inclusion and Ethics

Local researchers were included throughout the research process. Research protocols have been approved by local ethics review committees. No potential risk was involved in the current research, and all local researchers have discussed and approved the research protocol. All participants provided written informed consent, and local institutional review boards approved each study from all included cohorts. The analysis plan of the current study has been reviewed and approved by the Institutional Review Board of the Institute of Psychology, Chinese Academy of Sciences (No. H21102).

### Data Sharing

According to the success of the data sharing model of DIRECT Phase I data (REST-meta-MDD, http://rfmri.org/REST-meta-MDD), DIRECT Phase II data will also have 2 sharing stages. 1) Stage 1: coordinated sharing upon the publication of this announcing manuscript. To reduce conflict among the researchers, the consortium will review and coordinate the proposals submitted by interested researchers. The interested researchers first send a letter of intent to. Then, the consortium will send all the approved proposals to the applicant. The applicant should submit a new innovative proposal while avoiding conflict with approved proposals. The consortium would review and approve this proposal if there is no conflict. Once approved, this proposal would enter the pool of approved proposals and prevent future conflict. 2) Stage 2: unrestricted sharing after January 1, 2026. The researchers can perform any analyses of interest while not violating ethics. All codes have been made openly available (https://github.com/Chaogan-Yan/PaperScripts/tree/master/Chen_2023).

## Acknowledgments

This work was funded by the Sci-Tech Innovation 2030 – Major Project of Brain Science and Brain-inspired Intelligence Technology (No. 2021ZD0200600), the National Key R&D Program of China (No. 2017YFC1309902), the National Natural Science Foundation of China (No. 82122035, No. 81671774 and No. 81630031), the Key Research Program of the Chinese Academy of Sciences (No. ZDBS-SSW-JSC006), Beijing Nova Program of Science and Technology (No. Z191100001119104 and 20230484465), Beijing Natural Science Foundation (J230040), the Scientific Foundation of Institute of Psychology, Chinese Academy of Sciences (No. E2CX4425YZ and No. Y9CX422005), the China Postdoctoral Science Foundation (No. 2019M660847), the China National Postdoctoral Program for Innovative Talents (No. BX20200360), the Special Research Assistant Program of the Chinese Academy of Sciences (No. E2CX0624) and the Key R&D Program of Sichuan Province (No. 2023YFS0076). The support provided by the China Scholarship Council (CSC, No. 202104910248) during a visit of Xiao Chen to the Centre for Addiction and Mental Health is acknowledged. In addition, we would like to acknowledge the valuable insights provided by Prof. B.T. Thomas Yeo, whose comments helped to shape our understanding of the data and refine our analysis.

