## Supplementary_materials for "Subgenual Anterior Cingulate Cortex Functional Connectivity Abnormalities in Depression: Insights from Brain Imaging Big Data and Precision-Guided Personalized Intervention via Transcranial Magnetic Stimulation"

### Supplementary Information

### Sample Selection

We selected the final sample for the image analyses according to the following criteria for DIRECT Phase II data: (1) Bad image coverage. We excluded participants with imaging data that covered less than 90 percent of the group mask. The mask denoting the analysis area was generated by extracting all brain regions that 90% of all participants’ brain images covered. This discarded 15 participants. (2) Excessive head motion. Individuals with mean frame-wise displacement (FD) larger than 0.2 mm were excluded. Eighty-two individuals were discarded per this criterion (3) Missing demographical data. Next, we removed participants without necessary demographical information (i.e., age, sex, education, and head motion), which excluded 20 participants. This yielded a final sample containing 1574 MDD patients and 1308 HCs (Figure S6).

### Preprocessing

The preprocessing of fMRI data was conducted using the toolbox for Data Processing & Analysis for Brain Imaging on Surface V1.8_220915 (DPABISurf)^1^, which is based on fMRIPrep ^2^, FreeSurfer ^3^, ANTs ^4^, FSL ^5^, AFNI ^6^, SPM ^7^, PALM ^8^, dcm2niix ^9^, GNU Parallel ^10^, MATLAB R2020a (The MathWorks Inc., Natick, MA, US), Docker (https://docker.com) and DPABI V6.2_220915 ^11^. The preprocessing pipeline included: i) discarding the initial 10 time points; ii) converting data into BIDS format ^12^; iii) calling fMRIPrep 20.2.1 docker; iv) conducting anatomical data preprocessing as follows: correcting the T1-weighted image for intensity nonuniformity using N4BiasFieldCorrection ^13^; the corrected T1-weighted image was then skull-stripped with ANTs ^4^; fast (FSL 5.0.9)^14^ was used to segment brain tissue into cerebrospinal fluid (CSF), white-matter (WM) and gray-matter (GM); recon -all (FreeSurfer 6.0.1)^3^ was used to reconstruct brain surfaces; ANTs-derived brain masks and FreeSurfer-derived segmentations of the cortical GM were reconciled using a custom variation of the method of Mindboggle ^15^; Nonlinear registration with antsRegistration (ANTs 2.3.3) was performed for the volume-based spatial normalization to one standard space (MNI152NLin2009cAsym)^16^, using brain-extracted versions of both T1w reference and the T1w template; v) conducting functional data preprocessing as follows: generating a reference volume and its skull-stripped version using a custom methodology of fMRIPrep. The BOLD reference was then co-registered to the T1w reference using bbregister (FreeSurfer), which implements boundary-based registration ^17^. Co-registration was configured with six degrees of freedom. Head-motion parameters with respect to the BOLD reference (transformation matrices and six corresponding rotation and translation parameters) were estimated before any spatiotemporal filtering using mcflirt (FSL 5.0.9). BOLD runs were slice-time corrected using 3dTshift from AFNI 20160207 (RRID:SCR_005927)^6^. The BOLD time-series were resampled into standard space, generating a preprocessed BOLD run in MNI152NLin2009cAsym space; confounding time-series were calculated based on the preprocessed BOLD series in the original space: framewise displacement (FD), DVARS, and three region-wise global signals (extracted from the cerebrospinal fluid (CSF), the white matter (WM), and the whole-brain masks). FD was computed using Jenkinson (relative root mean square displacement between affines) and referred to as FD_Jenkinson_ ^5^. vi) Nuisance regression: The Friston 24-parameter model ^18^ was used to regress out head motion confounds. Additionally, mean framework displacement (FD) was used to address the residual effects of motion in group analyses^5^. Other sources of spurious variance (WM and CSF signals) were also removed from the data through linear regression to reduce respiratory and cardiac effects. Given that global signal regression (GSR) is an essential preprocessing step in identifying the TMS target ^19^, we performed preprocessing pipelines with and without GSR. Additionally, linear trends were included as a regressor to account for drifts in the blood oxygen level-dependent (BOLD) signal. vii) Finally, a bandpass temporal filter (0.01–0.1 Hz) was applied to the normalized functional images.

### Harmonization of site effects

ComBat models site-specific scaling factors by introducing terms for biological variables and scanners in a multivariate linear mixed effects regression. By employing empirical Bayesian criteria, it improves the estimation of studies with small sample sizes. ComBat assumes that the expected FC value can be modeled as a linear combination of batch effects and biological effects. Specifically, the error term in this model is modulated by additional site-specific scaling factors. Let y denote a given FC value:

$y =\alpha+ X^{T}\beta+\gamma+\delta\varepsilon$	(1)

where 𝛂 denotes the constant term; X^T^ denotes the design matrix for covariates of interest (diagnosis, age, sex, education, head motion); 𝛃 is a vector of regression coefficients corresponding to X; 𝛄 stands for addictive site effects (location parameter) while 𝛅 represents multiplicative site effects (scale parameter). Accordingly, harmonized FC values were defined as:

$y^{ComBat}= \frac{y-\hat{\alpha}-X^{T}\hat{\beta}-\hat{\gamma}}{\hat{\delta}}+\alpha+X^{T}\hat{\beta}$	(2)

where $\hat{\delta}$ and $\hat{\gamma}$ stand for the empirical Bayesian estimates of 𝛅 and 𝛄, respectively. y^ComBat^ was calculated with the “yw_Harmonization” function in DPABI ([http://rfmri.org/DPABI](http://rfmri.org/DPABISurf)). ComBat models biological and non-biological effects and algebraically removes the estimated additive and multiplicative batch effects. To empirically evaluate the capacity of ComBat in accounting for batch effects, we calculated the FCs between sgACC and two DLPFC clusters showing significant case-control differences using R-fMRI data with/without ComBat. A one-way ANOVA was performed to characterize the variability across all DIRECT sites (Figure S2-S3).

### The impact of GSR on group difference maps

Global signal regression (GSR) is the removal of the mean signal averaged across the entire brain via linear regression and is one of the most controversial preprocessing steps in resting state fMRI ^20,21^. On the one hand, GSR can increase the specificity of FC by efficiently reducing the impact of non-neuronal nuisance variables, such as head motion and respiration ^22-24^. On the other hand, GSR has been criticized for artificially shifting the distribution of the originally derived FCs and introducing spurious anti-correlations ^25,26^. Accordingly, not all previous studies implemented GSR, which might significantly alter case-control comparison results ^27^.

We demonstrated that GSR dramatically changed the group difference maps of sgACC FC profiles. One recent study examined the relationship between the sgACC-DLPFC anti-correlation and the TMS treatment outcomes using a large dataset (THREE-D) ^28^ and found that such correlation was largely carried by respiration-related signal patterns ^29^. Accordingly, most prior studies that intended to identify the TMS targets guided by sgACC-related FCs tend to implement GSR to remove non-neural noises as much as possible. In contrast, other studies delineating general case-control differences preferred not to implement GSR. Here, we showed that GSR might induce notable alterations in case-control differences of FC maps ^27^. Thus, we also provided FC maps without GSR in the supplementary materials for those interested in information regarding the neuropathological basis of MDD without GSR.

#### **Identification of individualized TMS targets by dual regression and seed map approach**

After obtaining the reliable statistical maps from the DIRECT sample, we used the dual regression approach and the seed map approach to identify individualized TMS targets in the SID-TMS and the CUD-TMS datasets.

In the DR approach, first, a group-level statistical spatial map was used as a spatial regressor in the general linear model (GLM) to identify the temporal dynamic of the group-level map (similar to spatial correlation with this group-level statistical spatial map). A time series associated with the group-level statistical spatial map was generated. This time course measured how the group-level template fit into the brain BOLD signal spatial pattern at every time point. We used three group-level sgACC-FC statistical maps (i.e., the group difference map, the mean MDD sgACC-FC map, and the mean HC sgACC-FC map) as group templates in the DR-based algorithm. As the two clusters that survived FDR correction (*q* < 0.05) were too small to be used as group templates in the DR computation, we expanded them by applying a more liberal multiple comparison correction threshold (FDR *q* < 0.1). In addition, we reversed the mean MDD/HC sgACC-FC maps to generate the most sgACC-anticorrelated brain area in DLPFC before conducting regressions.

Second, the time course derived in the first step was used as a temporal regressor to estimate subject-specific spatial maps (similar to temporal correlation with the previous time course). These subject-specific spatial maps were considered the best approximation of the group-level template for single participants. Subsequently, we applied a threshold to the subject-specific spatial maps, retaining only the top-n voxels within the DLPFC mask. The final coordinates for the individualized TMS targets were defined as the centroids of the largest clusters within the DLPFC region on the thresholded individual-level spatial maps. The thresholds were related to the type and size of the group template input into the DR algorithm (Figure S9). We chose BA 46 as the DLPFC mask in the DR approach because prior studies have reported that BA 46 is more anticorrelated to sgACC than alternative DLPFC definitions (35) (e.g., BA 9, Beam F3, or the 5 cm method) and suggested that a more anterior and lateral location was related to better treatment response in MDD TMS treatment (7, 37). Considering that the entire DR approach is conducted within the DLPFC area, we used data that had not undergone GSR during preprocessing to calculate TMS targets in order to avoid the influence of signals from unrelated brain regions.

Additionally, we calculated the individualized target coordinates using the seed map approach, following the methods described by Fox et al. and Cash et al. The image preprocessing procedure is the same as that described by Fox et al. ^19^ The seed map is defined as all gray matter areas except for the DLPFC region. This DLPFC region is defined as the union of four 20 mm radius spheres centered at the centroids of BA46 (MNI [-44,40,29]), BA9 (MNI [-36,39,43]), the 5cm method group target (MNI [-41,16,54]), and the F3 group target (MNI [-37,26,49]) following the methodology described by Cash et al. ^30^. Pearson's correlation coefficients were computed between this extracted time series and those of all other DLPFC voxels. The voxel-wise threshold was set to *t* = -3 to meet the requirement that the final target’s cluster size be greater than 1. The final TMS target was defined as the most negatively functionally connected cluster in the DLPFC area.

### Tables

Table S1. DIRECT site info, sample sizes and previous publications based on the shared data.

| Serial Number | Sites (cohorts) | Principal investigators | Data organizer | N | | Publications |
| --- | --- | --- | --- | --- | --- | --- |
|  |  |  |  | MDD | NC |  |
| 1 | Faculty of Psychology, Southwest University | Jiang Qiu | Xin-Ran Wu | 282 | 251 | Cheng et al., 2016/ Ye et al., 2015 ^31,32^ |
| 2 | Department of Psychiatry, The First Affiliated Hospital of Chongqing Medical University | Li Kuang | Qian Huang | 151 | 63 | Cao et al., 2016 ^33^ |
| 3 | National Clinical Research Center for Mental Disorders (Peking University Sixth Hospital) & Key Laboratory of Mental Health, Ministry of Health (Peking University) | Tian-Mei Si | Yan-Kun Wu | 20 | 20 | Wang et al., 2013/Wang et al., 2015 ^34,35^ |
| 4 | Mental Health Center, West China Hospital, Sichuan University | Tao Li | Hua Yu | 34 | 30 | Yang et al., 2015 ^36^ |
| 5 | First Hospital of Shanxi Medical University | Ke-Rang Zhang | Ai-Xia Zhang | 139 | 63 | Li et al., 2014 ^37^ |
| 6 | The Institute of Mental Health, Second Xiangya Hospital of Central South University | Zhe-Ning Liu | Yi-Cheng Long | 67 | 106 | Zeng et al., 2020/Tan et al., 2021/Long et al., 2020 ^38-40^ |
| 7 | Affiliated Brain Hospital of Guangzhou Medical University | Li-Ping Cao | Jian-Shan Chen | 24 | 30 | Cheng et al., 2021 ^41^ |
| 8 | The First Affiliated Hospital of Xi’an Jiaotong University, Xi’an Central Hospital | Jian Yang / Xiao-Ping Wu | Peng-Feng Sun | 47 | 42 | Wu et al., 2016 ^42^ |
| 9 | Department of Psychosomatics and Psychiatry, Zhongda Hospital, School of Medicine, Southeast University | Yong-Gui Yuan | Xiao-Yun Liu | 47 | 26 | Hou et al., 2018/Hou et al., 2018 ^43,44^ |
| 10 | Beijing Anding Hospital, Capital Medical University | Chuan-Yue Wang | Feng Li | 85 | 70 | Zheng et al., 2018/Liu et al., 2017 ^45,46^ |
| 11 | Department of Neurology, Affiliated ZhongDa Hospital of Southeast University | Chun-Ming Xie | Can-Can He | 116 | 68 | Yuan et al., 2008 ^47^ |
| 12 | The Second Xiangya Hospital of Central South University | Guang-Rong Xie | Xi-Long Cui | 63 | 32 | Yang et al., 2017 ^48^ |
| 13 | Department of Clinical Psychology, Suzhou Suzhou Psychiatric Hospital, The Affiliated Guangji Hospital of Soochow University | Yan-Song Liu | Feng-Nan Jia | 33 | 29 | N/A |
| 14 | The First Affiliated Hospital of Anhui Medical University | Yong-Qiang Yu | Jia-Jia Zhu | 139 | 147 | Zhu et al., 2020/Zhao et al., 2022 ^49,50^ |
| 15 | Xiangya 2nd Hospital, Central South University | Xiang Wang | Xin-Lei Ji | 35 | 35 | N/A |
| 16 | The First Affiliated Hospital of Jinan University | Ying Wang | Guan-Mao Chen | 83 | 81 | N/A |
| 17 | First Affiliated Hospital of Kunming Medical University | Xiu-Feng Xu / Yu-Qi Cheng | Zhao-Song Chu | 47 | 50 | He et al., 2022 ^51^ |
| 18 | Huaxi MR Research Center, West China Hospital of Sichuan University | Qi-Yong Gong | Tao-Lin Chen | 34 | 34 | Chen et al., 2017 ^52^ |
| 19 | The Second Xiangya Hospital of Central South University | Wen-Bin Guo/Jing-Ping Zhao | Xiao-Xiao Shan | 32 | 32 | Guo et al., 2014/Guo et al., 2018 ^53,54^ |
| 20 | The First Affiliated Hospital, College of Medicine,Zhejiang University | Yang Hong | Tao Chen | 20 | 20 | N/A |
| 21 | Xijing Hospital | Hua-Ning Wang/Bao-Juan Li | Bao-Juan Li | 63 | 56 | N/A |
| 22 | West China Hospital, Sichuan University | Wei Zhang | Min-Lan Yuan | 35 | 25 | Xiao et al., 2021 ^55^ |
| 23 | Shenzhen Mental Health Center, Shenzhen Kangning Hospital | Jian-ping Lu | Zhen-Peng Xue | 64 | 31 | N/A |
|  |  |  |  | 1660 | 1341 |  |

Table S2. DIRECT image acquisition by site.

| Serial Number | Scanner | Receive coil | TR (ms) | TE (ms) | Flip Angle (∘) | Thickness/gap (mm) | Slice number | Time points | Voxel size (mm) | FOV | Eyes open vs. closed | Slice order |
| --- | --- | --- | --- | --- | --- | --- | --- | --- | --- | --- | --- | --- |
| 1 | Siemens Tim Trio 3T | 12-channel | 2000 | 30 | 90 | 3/1 | 32 | 242 | 3.44 × 3.44 × 4.00 | 220 × 220 | closed | Interleaved Descending |
| 2 | GE Signa 3T | 8-channel | 2000 | 40 | 90 | 4/0 | 33 | 240 | 3.75 × 3.75 × 4.00 | 240 × 240 | closed | Interleaved Ascending |
| 3 | Siemens Tim Trio 3T | 32-channel | 2000 | 30 | 90 | 4.0/0.8 | 30 | 210 | 3.28 × 3.28 × 4.80 | 210 × 210 | closed | Interleaved Descending |
| 4 | Philips Achieva 3T TX | 8-channal | 2000 | 30 | 90 | 4.0/0 | 38 | 240 | 3.75 × 3.75 × 4.00 | 240 × 240 | closed | Interleaved Ascending |
| 5 | Siemens Tim Trio 3T | 32-channel | 2000 | 30 | 90 | 3.00/1.52 | 32 | 212 | 3.75 × 3.75 × 4.52 | 240 × 240 | closed | Interleaved Descending |
| 6 | Philips Gyroscan Achieva 3T | 32-channel | 2000 | 30 | 90 | 4/0 | 36 | 250 | 1.67 × 1.67 × 4.00 | 240 × 240 | closed | Interleaved Ascending |
| 7 | Philips Achieva X-series 3T | 8-channel | 2000 | 30 | 90 | 4.0/0.6 | 33 | 240 | 3.44 × 3.44 × 4.60 | 220 × 220 | closed | Interleaved Ascending |
| 8 | GE Excite 1.5T | 16-channel | 2500 | 35 | 90 | 4/0 | 36 | 150 | 4.00 × 4.00 × 4.00 | 256 × 256 | closed | Interleaved Ascending |
| 9 | Siemens Verio 3T | 12-channel | 2000 | 25 | 90 | 4/0 | 36 | 240 | 3.75 × 3.75 × 4.00 | 240 × 240 | closed | Interleaved Descending |
| 10 | Siemens Tim Trio 3T | 32-channel | 2000 | 30 | 90 | 3.5/0.7 | 33 | 240 | 3.12 × 3.12 × 4.20 | 200 × 200 | closed | Interleaved Descending |
| 11 | Siemens Verio 3T | 12-channel | 2000 | 25 | 90 | 4/0 | 36 | 240 | 3.75 × 3.75 × 4.00 | 240 × 240 | closed | Interleaved Descending |
| 12 | Siemens Tim Trio 3T | 32-channel | 2500 | 25 | 90 | 3.5/0 | 39 | 200 | 3.75 × 3.75 × 3.50 | 240 × 240 | closed | Interleaved Descending |
| 13 | Siemens Skyra 3T | 32-channel | 2000 | 30 | 90 | 3.5/0 | 32 | 240 | 3.50 × 3.50 × 3.50 | 224 × 224 | closed | Interleaved Ascending |
| 14 | GE Discovery MR750w 3T | 24-channel | 2000 | 30 | 90 | 3/1 | 35 | 185 | 3.40 × 3.40 × 4.00 | 220 × 220 | closed | Interleaved Ascending |
| 15 | Siemens Skyra 3T X | 32-channel | 2000 | 30 | 90 | 4/0 | 36 | 240 | 3.44 × 3.44 × 4.00 | 220 × 220 | open | Interleaved ascending |
| 16 | GE Discovery MR750 3T | 8-channel | 2000 | 25 | 90 | 3/1 | 35 | 200 | 3.75 × 3.75 × 4.00 | 240 × 240 | closed | Interleaved Descending |
| 17 | Philips Achieva 3T | 16-channel | 4400 | 35 | 90 | 3 | 50 | 240 | 1.80 × 1.80 × 3.00 | 230 × 230 | closed | Interleaved Ascending |
| 18 | GE Signa 3T | 8-channel | 2000 | 30 | 90 | 5/0 | 30 | 200 | 3.75 × 3.75 × 5.00 | 240 × 240 | closed | Interleaved Ascending |
| 19 | Siemens Skyra 3T | 32-channel | 2000 | 30 | 90 | 4.0/0.6 | 33 | 240 | 3.40 × 3.40 × 4.60 | 220 × 220 | closed | Interleaved Descending |
| 20 | GE Discovery 3T | 16-channel | 2000 | 30 | 90 | 3.2/0.4 | 43 | 200 | 3.40 × 3.40 × 3.40 | 220 × 220 | closed | Interleaved Descending |
| 21 | GE Discovery MR750 3T | 32-channel | 2000 | 30 | 90 | 3.5/0 | 45 | 210 | 3.75 × 3.75 × 3.50 | 240 × 240 | closed | Interleaved Ascending |
| 22 | Philips Achieva 3T | 8-channel | 2000 | 30 | 90 | 4/0 | 38 | 240 | 3.75 × 3.75 × 4.00 | 240 ×240 | closed | Interleaved Descending |
| 23 | GE MR750 3T | 8-channel | 2000 | 30 | 90 | 3.2/0 | 43 | 240 | 3.4375 × 3.4375 × 3.2 | 220 × 220 | closed | Interleaved Descending |

**Figures**

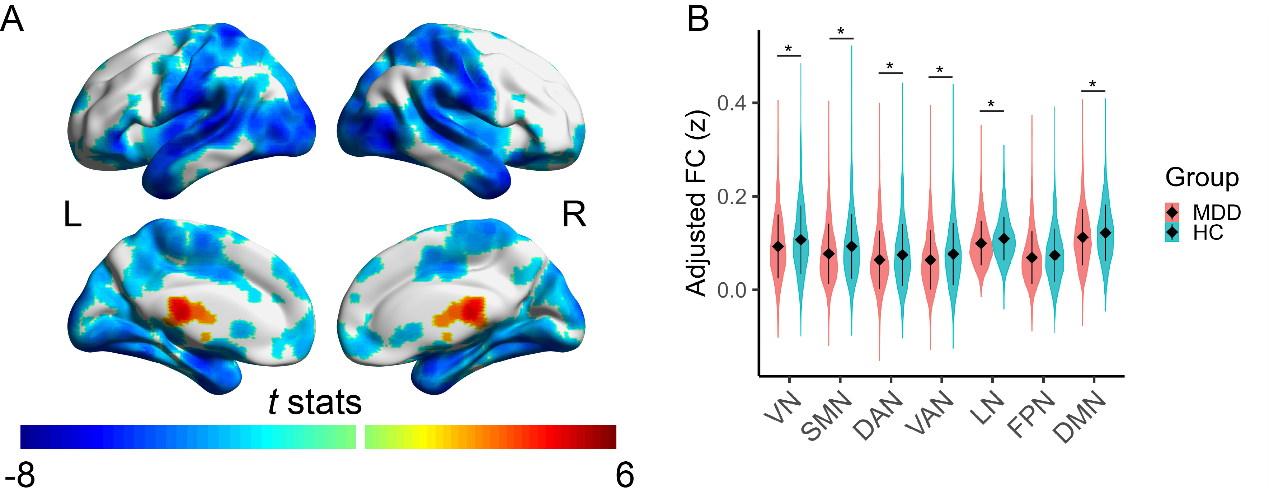

Figure S1. Case-control differences of subgenual anterior cingulate cortex (sgACC) FC profiles when global signal regression (GSR) was not implemented. (A) The FDR corrected (*q* < 0.05) brain render of the group difference map without GSR. (B) Group differences of FCs between sgACC and visual network (VN), somatomotor network (SMN), dorsal attention network (DAN), ventral attention network (VAN), limbic network (LN), frontoparietal network (FPN), and default mode network (DMN). *: significant after Bonferroni correction. Abbreviations: L, left hemisphere; R, right hemisphere.

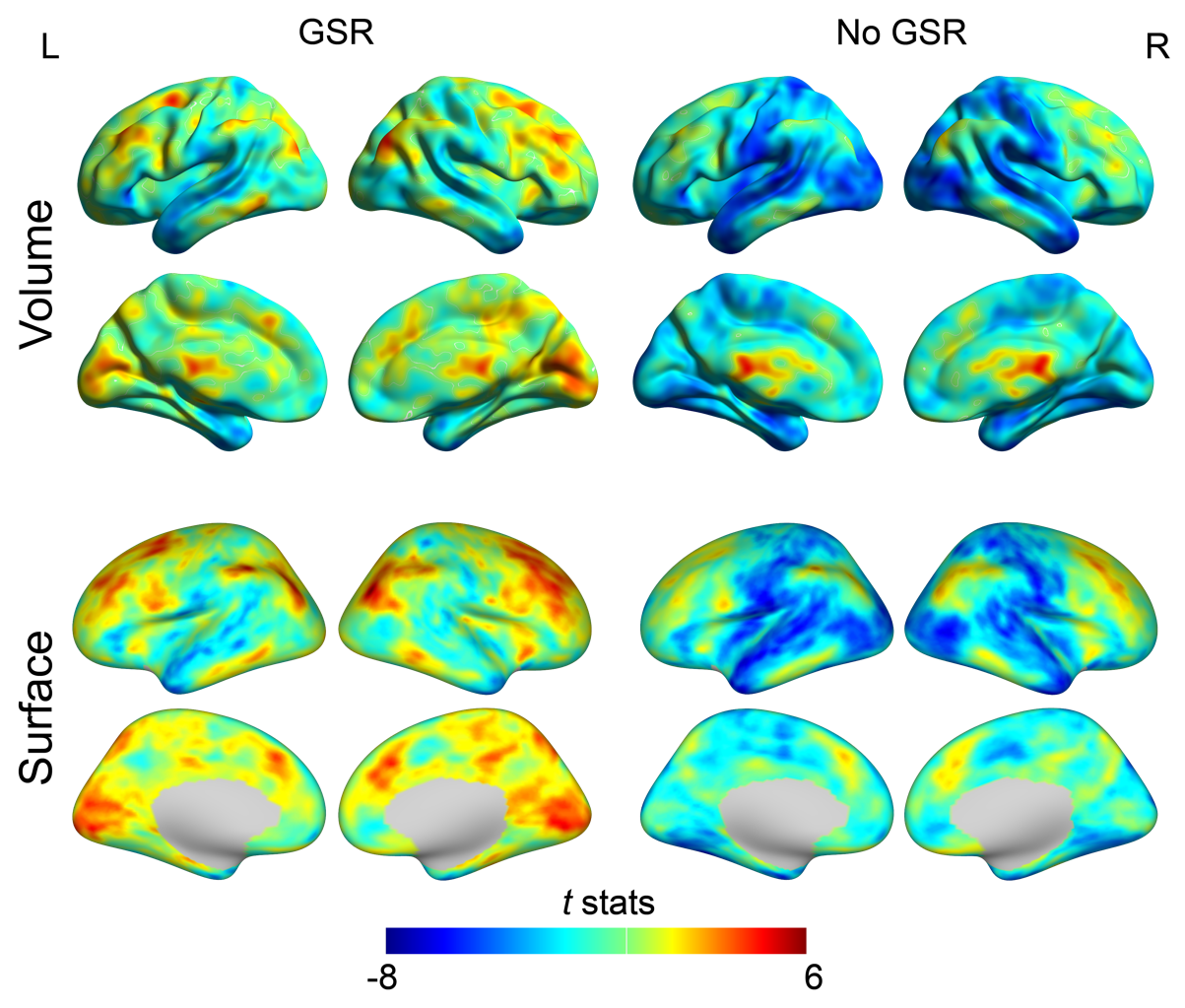

Figure S2. Uncorrected group difference *t* statistic maps with/without global signal regression (GSR) as one of the preprocessing steps. The upper panel shows the t statistic maps calculated in the volume space, and the lower panel shows the t statistic maps in the surface space. Note the general alignment between the results in the volume/surface spaces. Abbreviations: L, left hemisphere; R, right hemisphere.

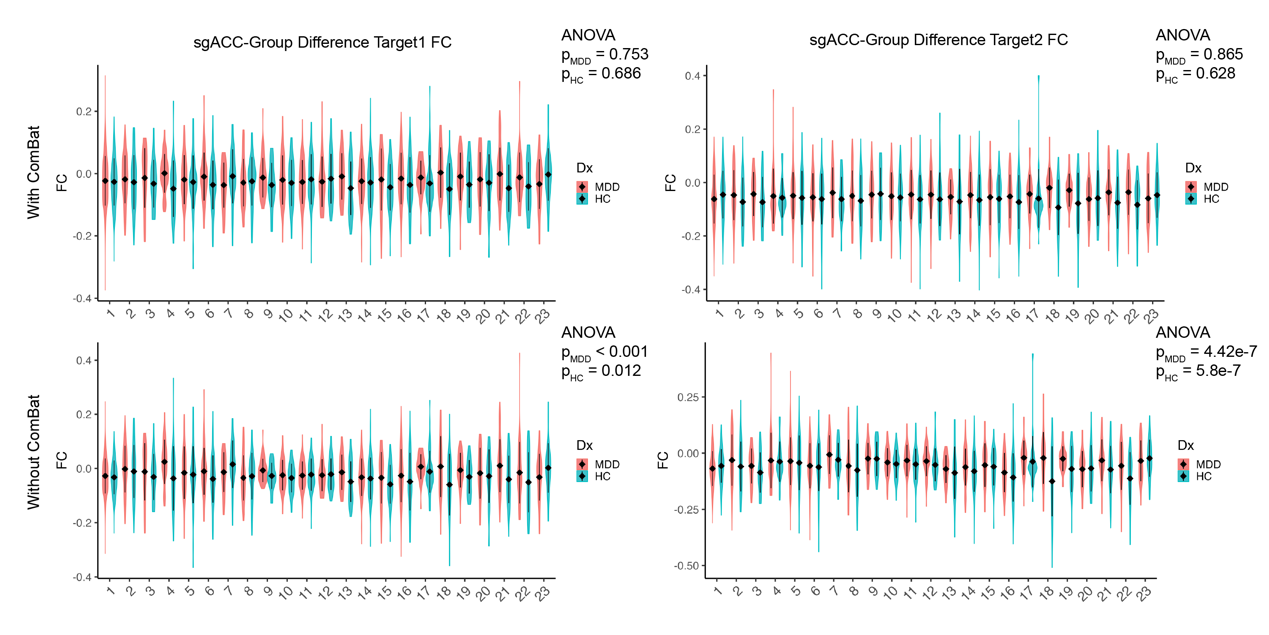

Figure S3. Raw FC data with global signal regression (GSR) for MDD and HC subjects from the DIRECT dataset. The top row shows FC values between sgACC and two group difference targets normalized using comBat. The bottom row shows raw FC values without comBat. ANOVA p-values refer to one-way analyses of variance across sites for FC values. GSR was included in the preprocessing pipeline.

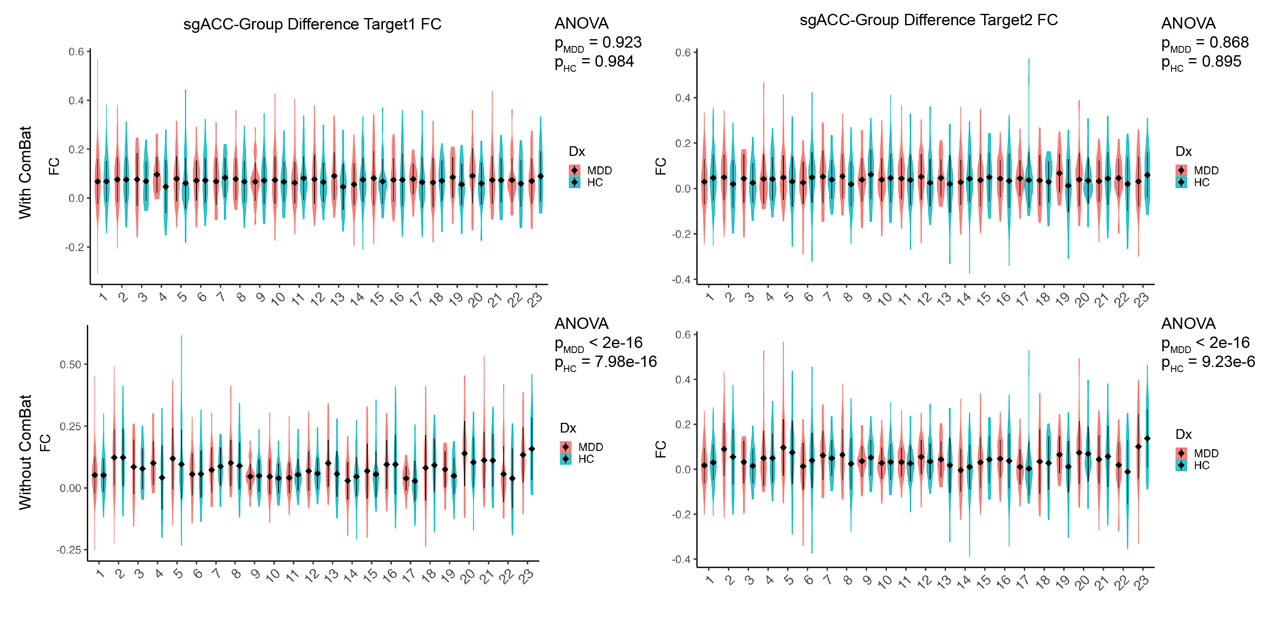

Figure S4. Raw FC data without global signal regression (GSR) for MDD and HC subjects from the DIRECT dataset. The top row shows FC values between sgACC and two group difference targets normalized using comBat. The bottom row shows raw FC values without comBat. ANOVA p-values refer to one-way analyses of variance across sites for FC values. GSR was not included in the preprocessing pipeline.

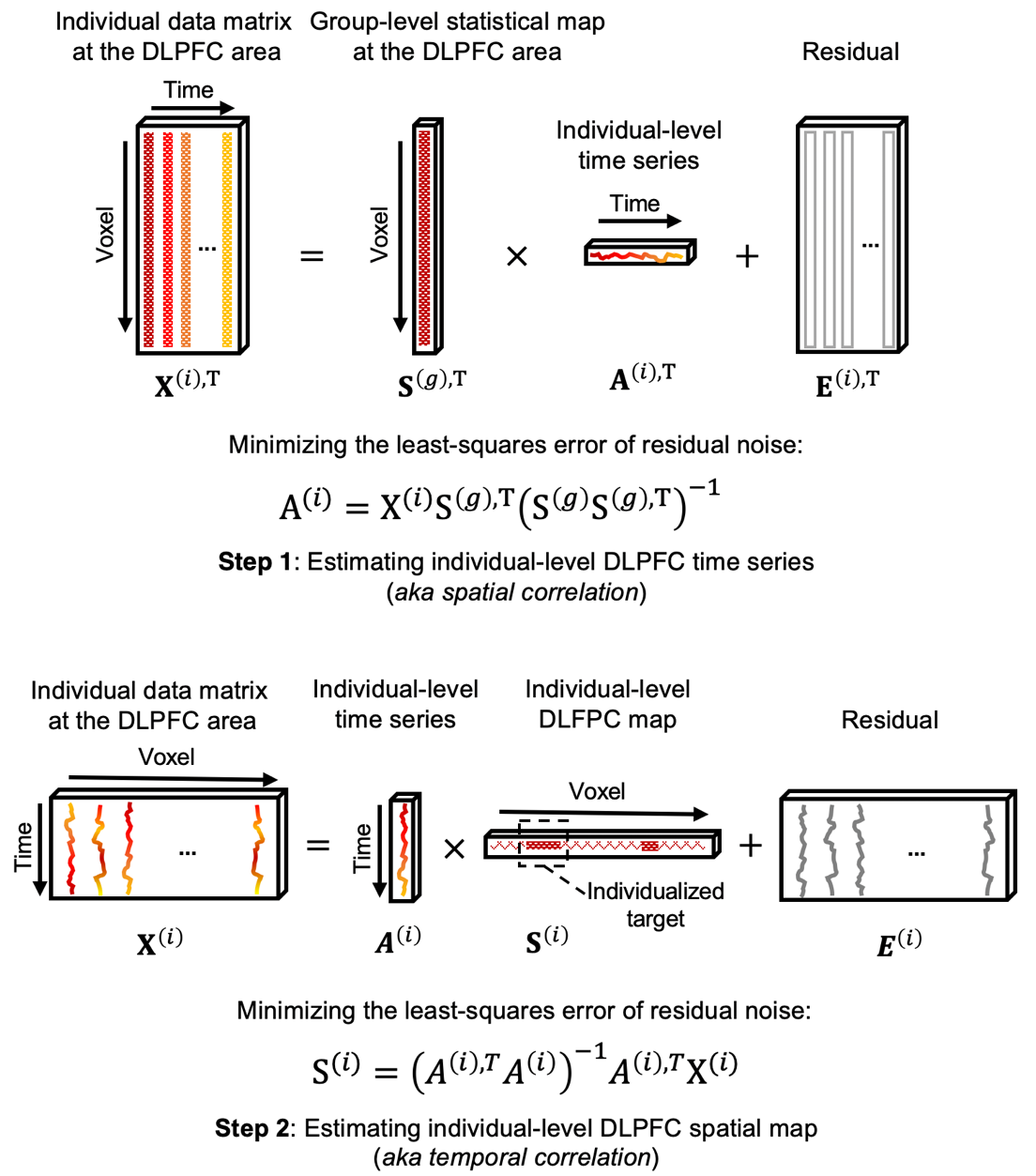

Figure S5. Schematic of the dual regression based individual TMS target identification. In the present study, $X^{(i)}$represents the R-fMRI data matrix of an individual patient. $S^{\left( g \right)}$ represents a group-level spatial template, such as the sgACC-FC group difference map. $A^{(i)}$ represents the estimated individual-level time series for the group template. $S^{(i)}$ represents the estimated individual-level spatial map used for TMS targeting.

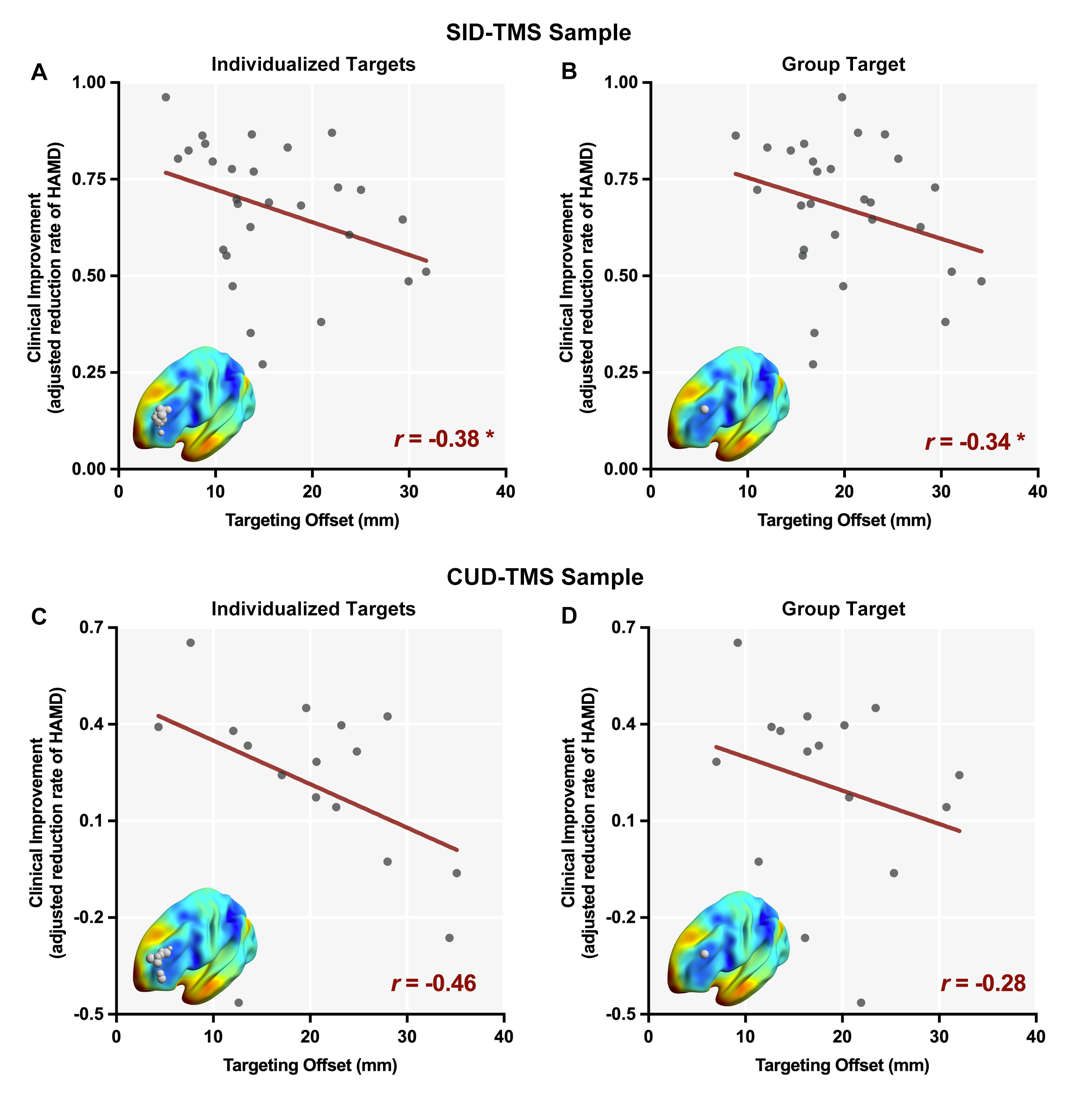

Figure S6. Individualized targets derived from the mean HC sgACC-FC map using the DR-based algorithm exhibited greater clinical efficacy than the corresponding group targets. Clinical efficacy was characterized by computing the correlations between the offset distances of TMS targets and the clinical improvements observed in the SID-TMS and CUD-TMS datasets. The TMS targeting offset distance was defined as the Euclidean distance between the actual rTMS stimulation coordinates and the individualized or group targets. Clinical improvement was defined as the HAMD reduction during rTMS treatment, adjusted for age, sex, and head motion. The locations of the targets are displayed on the cortex. The sizes of the spheres indicate the magnitudes of reductions in the Hamilton Depression Rating Scale (HAMD). (A-B) The clinical efficacy of the individualized and group targets in the SID-TMS sample. (C-D) The clinical efficacy of the individualized and group targets in the CUD-TMS sample. Abbreviations: HC, healthy control; sgACC-FC: functional connectivity using the subgenual anterior cingulate cortex as the seed. **p* < 0.05.

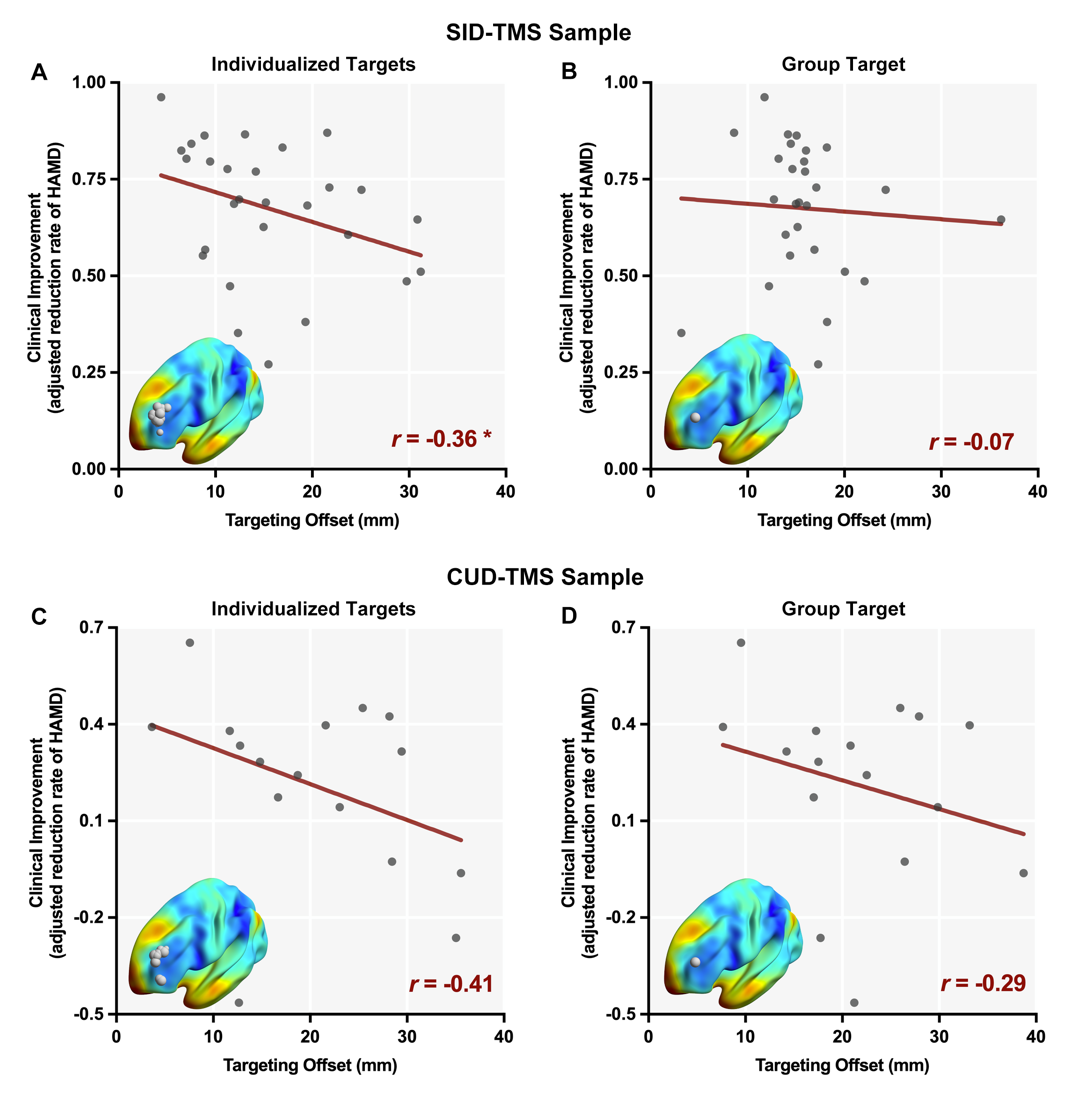

Figure S7. Individualized targets derived from the mean MDD sgACC-FC map exhibited greater clinical efficacy than the corresponding group targets. Clinical efficacy was characterized by computing the correlations between the offset distances of TMS targets and the clinical improvements observed in the SID-TMS and CUD-TMS datasets. The TMS targeting offset distance was defined as the Euclidean distance between the actual rTMS stimulation coordinates and the individualized or group targets. Clinical improvement was defined as the HAMD reduction during rTMS treatment, adjusted for age, sex, and head motion. The locations of the targets are displayed on the cortex. The sizes of the spheres indicate the magnitudes of reductions in the Hamilton Depression Rating Scale (HAMD). (A-B) The clinical efficacy of the individualized and group targets in the SID-TMS sample. (C-D) The clinical efficacy of the individualized and group targets in the CUD-TMS sample. Abbreviations: MDD, major depressive disorder; sgACC-FC: functional connectivity using the subgenual anterior cingulate cortex as the seed. *p < 0.05.

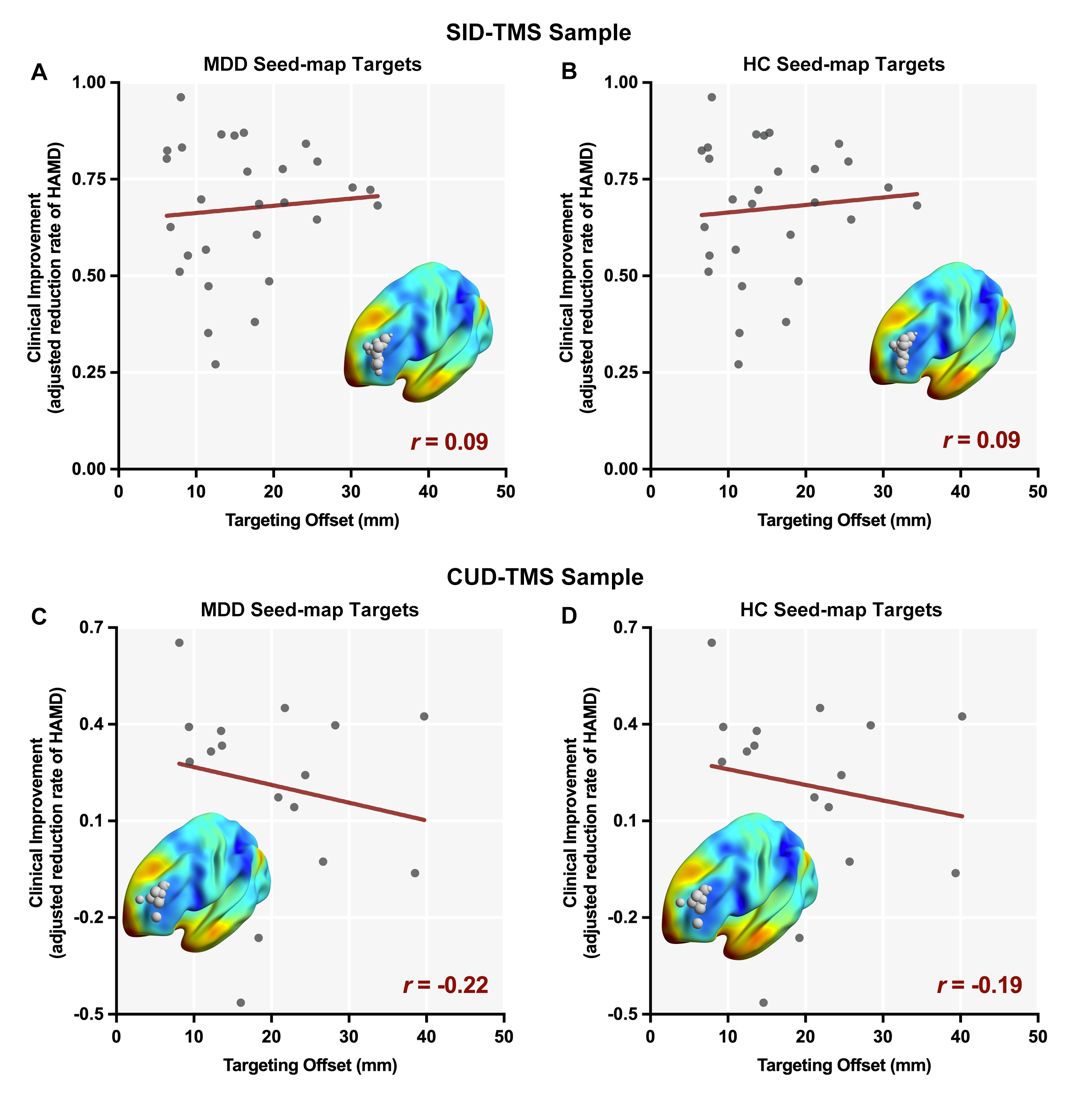

Figure S8. Clinical significance of the Individualized targets derived from the seed-map approach. The mean MDD/HC sgACC-FC map, excluding the DLPFC region, was used as the seed map. Clinical efficacy was characterized by computing the correlations between the offset distances of TMS targets and the clinical improvements observed in the SID-TMS and CUD-TMS datasets. The TMS targeting offset distance was defined as the Euclidean distance between the actual rTMS stimulation coordinates and the individualized or group targets. Clinical improvement was defined as the HAMD reduction during rTMS treatment, adjusted for age, sex, and head motion. The locations of the targets are displayed on the cortex. The sizes of the spheres indicate the magnitudes of reductions in the Hamilton Depression Rating Scale (HAMD). (A-B) The clinical efficacy of the individualized and group targets in the SID-TMS sample. (C-D) The clinical efficacy of the individualized and group targets in the CUD-TMS sample. Abbreviations: DLPFC, the left dorsal lateral prefrontal cortex; HC, healthy control; MDD, major depressive disorder; sgACC-FC: functional connectivity using the subgenual anterior cingulate cortex as the seed.

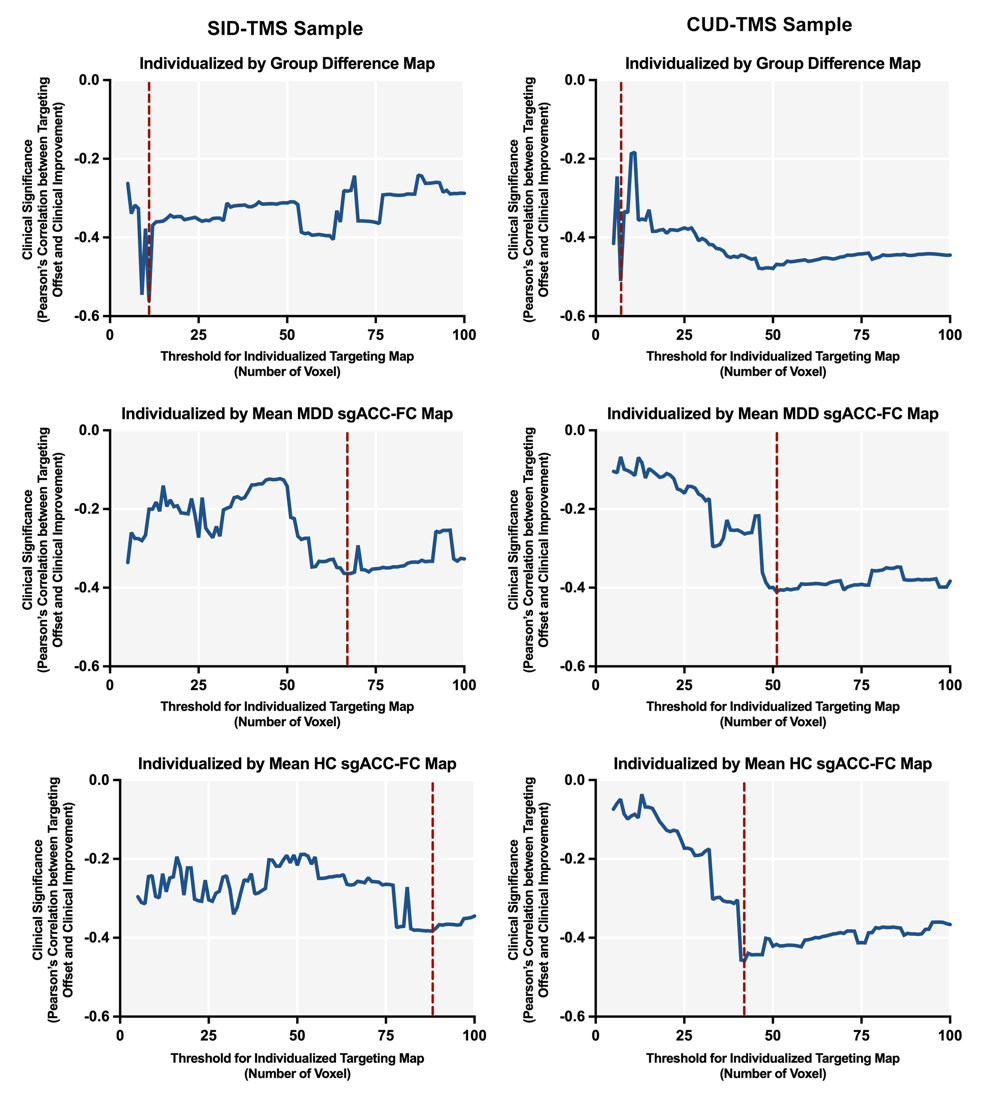

Figure S9. Variation of clinical significance at different thresholds of cluster size for the targets derived from the dual regression (DR)-based individualizing approach. Clinical significance was assessed by computing the correlations between the offset distances of TMS targeting and the clinical improvements observed in the patients (e.g., the lower the negative correlation value, the better the clinical significance of the target). The red line represents the cluster size threshold for targets applied in the main text.

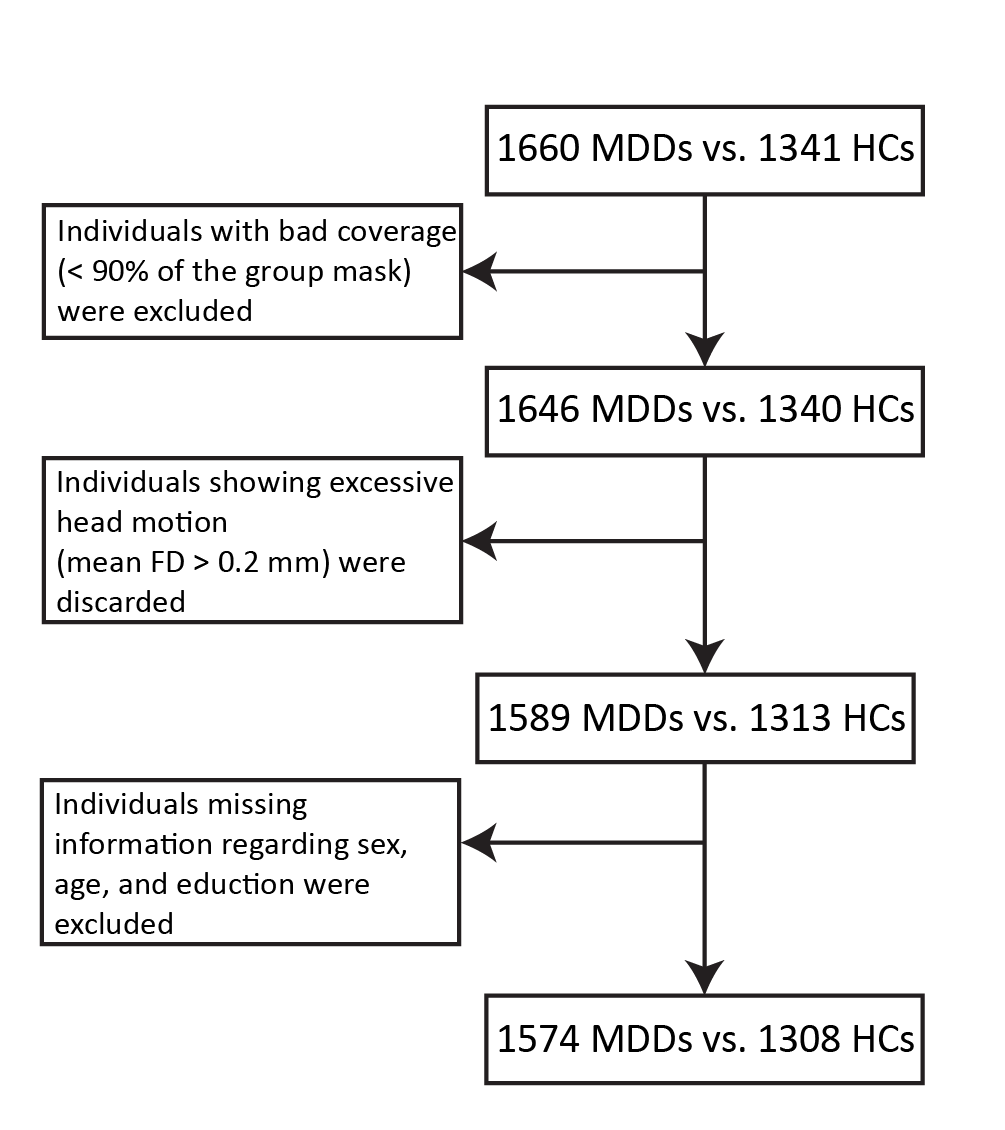

Figure S10. Flowchart depicting the sample selection procedure.

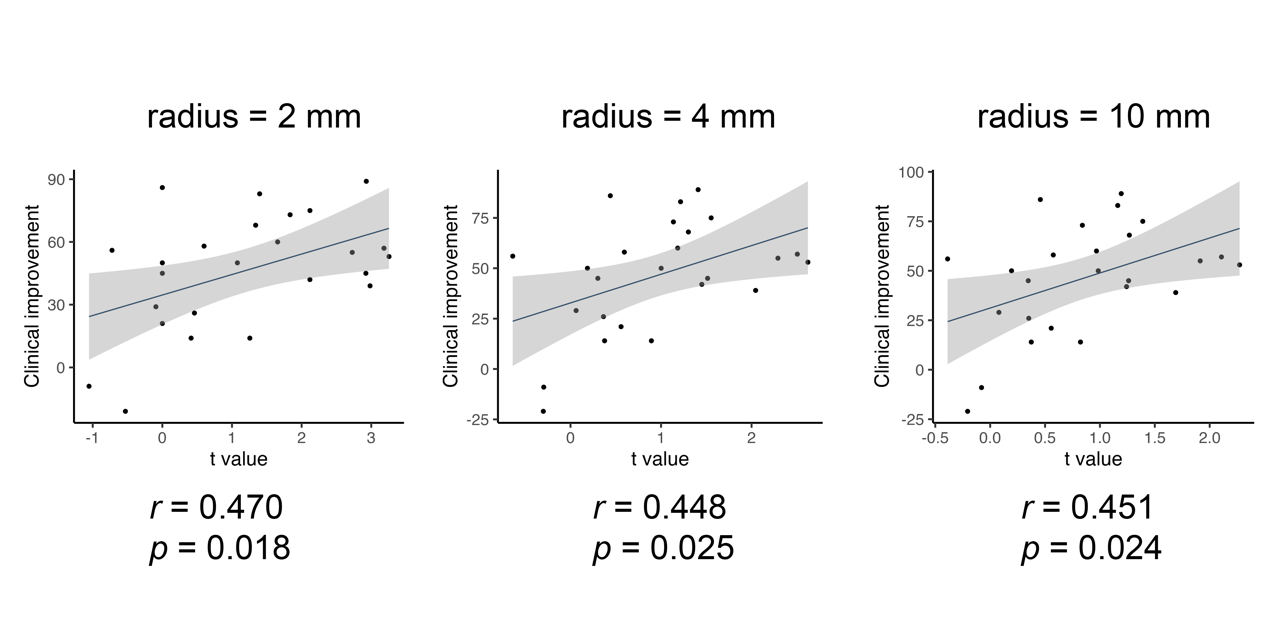

Figure S11. The relationship between TMS clinical efficiency and targets’ group differences extracted using spheres of different radiuses.
